# Assessment of current taxonomic assignment strategies for metabarcoding eukaryotes

**DOI:** 10.1101/2020.07.21.214270

**Authors:** Jose S. Hleap, Joanne E. Littlefair, Dirk Steinke, Paul D. N. Hebert, Melania E. Cristescu

## Abstract

The effective use of metabarcoding in biodiversity science has brought important analytical challenges due to the need to generate accurate taxonomic assignments. The assignment of sequences to a generic or species level is critical for biodiversity surveys and biomonitoring, but it is particularly challenging. Researchers must select the approach that best recovers information on species composition. This study evaluates the performance and accuracy of seven methods in recovering the species composition of mock communities which vary in species number and specimen abundance, while holding upstream molecular and bioinformatic variables constant. It also evaluates the impact of parameter optimization on the quality of the predictions. Despite the general belief that BLAST top hit underperforms newer methods, our results indicate that it competes well with more complex approaches if optimized for the mock community under study. For example, the two machine learning methods that were benchmarked proved more sensitive to the reference database heterogeneity and completeness than methods based on sequence similarity. The accuracy of assignments was impacted by both species and specimen counts which will influence the selection of appropriate software. We urge the usage of realistic mock communities to allow optimization of parameters, regardless of the taxonomic assignment method used.

## BACKGROUND

Accurate taxonomic classification of sequences recovered through metabarcoding is essential to ascertain the species encountered in biodiversity surveys (Bazinet & Cummings 2012; Mizrahi-Man *et al*. 2013; Richardson *et al*. 2017; Fosso *et al*. 2018; Heeger *et al*. 2018), biomonitoring (Bazinet & Cummings 2012; Mizrahi-Man *et al*. 2013; Richardson *et al*. 2017; Fosso *et al*. 2018; Heeger *et al*. 2018), and to detect invasive species (Guo *et al*. 2010; Gillet *et al*. 2018). Over the past five years, several algorithms have been developed to address the need for reliable, consistent taxonomic assignments (Figure 1). These methods employ diverse approaches: direct comparison of local alignments (e.g. BLAST; Altschul *et al*. 1997), post-processing of local alignments (e.g. MEGAN-like Last Common Ancestor LCA algorithms; (Clemente *et al*. 2011; Mitra *et al*. 2011; Wood & Salzberg 2014; Kahlke & Ralph 2019), machine learning techniques based on k-mers (Rosen *et al*. 2011; Lan *et al*. 2012; Murali *et al*. 2018), phylogenetic techniques (Munch *et al*. 2008; Nguyen *et al*. 2014; Janssen *et al*. 2018; Zheng *et al*. 2018), and probabilistic methods (Somervuo *et al*. 2016; Axtner et al. 2019).

**Figure 1.**
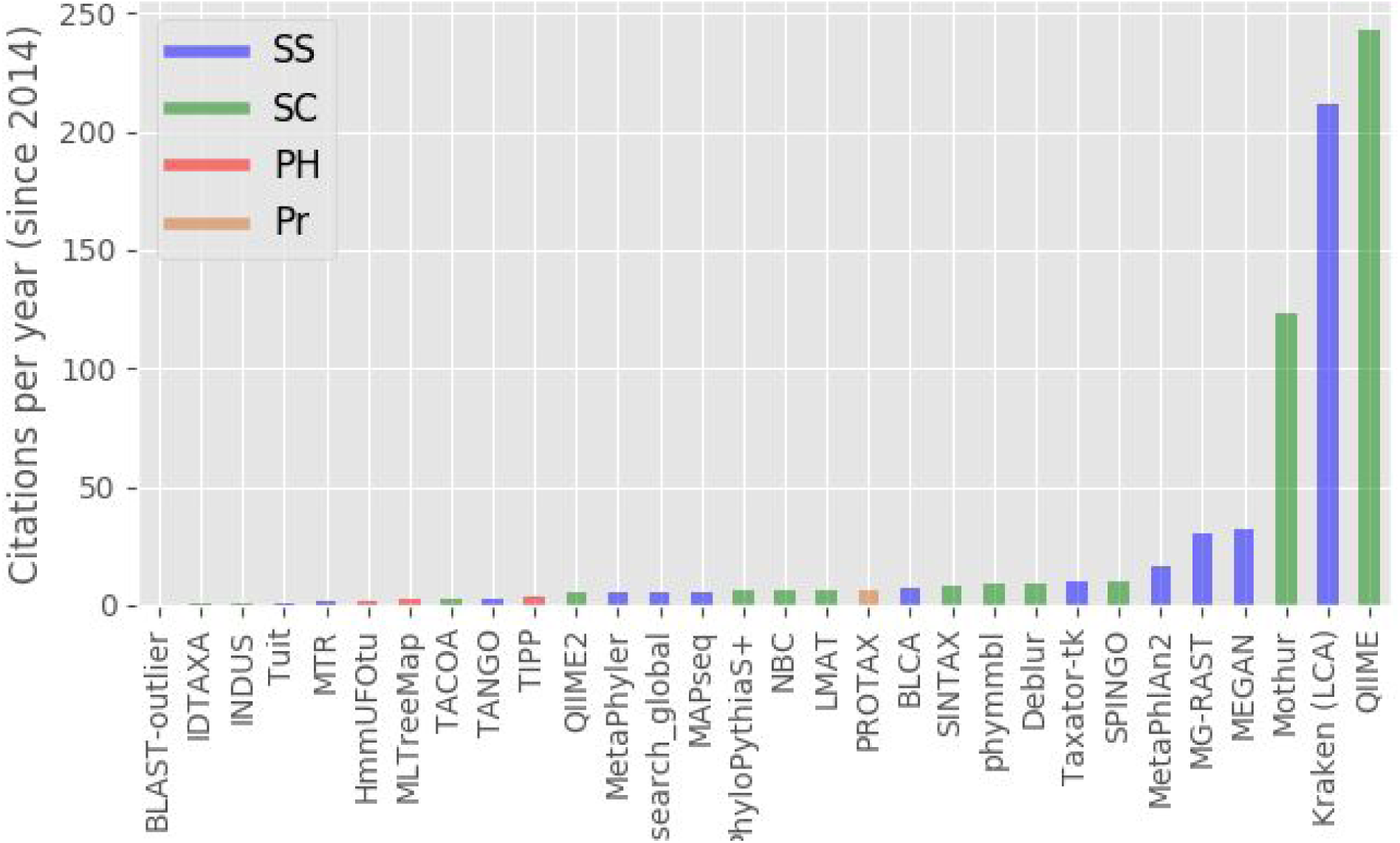
Annual citation rates over the last five years for 33 taxonomic assignment methods. Each method is assigned to one of four major categories - SC: Sequence composition, SS: Sequence Similarity, PH: Phylogenetic, PR: Probabilistic. Citation counts derive from a Google Scholar for the term “taxonomic assignment”, filtered by year.

Bazinet and Cummings (2012) proposed a system that classified the main supervised learning approaches. We now extend this categorization system by classifying taxonomic assignment programs into four major strategies (Figure 2, Table 1): sequence similarity (SS), sequence composition (SC), phylogenetics (PH, also referred to as model-based by Richardson *et al*. 2017), and probabilistic (PR). All SS methods use global or local alignments to directly compare query sequences to sequences in a reference database. BLAST (Altschul *et al*. 1997) is the best known of these methods. Methods based on SC classify sequences by extracting compositional features (e.g. nucleotide frequency patterns) before building a model that links these profiles to specific taxonomic groups. The Ribosomal Database Project (RDP; Naive Bayes-based) is the most widely used SC classifier (Wang *et al*. 2007). PH methods rely on the phylogenetic placement of a sequence; they utilize a tree reconstruction method, such as Maximum Likelihood, to obtain a *de novo* phylogeny or perform a read recruitment strategy against an existing reference tree (Janssen *et al*. 2018) using an evolutionary placement algorithm (EPA; Barbera *et al*. 2019). Finally, PR methods assess the probability of correctly placing a query sequence to a particular taxonomic level by employing a probabilistic framework such as multinomial regression (Figure 2; Somervuo *et al*. 2016).

**Table 1.** Strengths and weaknesses of the main methods for taxonomic assignment. Reliability refers to how often expected results are recovered; Availability depicts the ease of obtaining and installing the program (including current support); Scalability is the capacity to upsize the test; Understandability is the ease of comprehension of the algorithm by non-technical users; Ease of use refers to how easy is to install and run the program.

|  | Sequence<br>Similarity (SS) | Sequence<br>Composition<br>(SC) | Phylogenetic<br>(Ph) | Probabilistic<br>(Pr) |
| --- | --- | --- | --- | --- |
| **Reliability** | High/Low* | High/Low* | High*** | High |
| **Availability** | High | Medium | Medium | Low |
| **Scalability** | Medium | High/Low** | Very low | High |
| **Understandability** | High | Low | High | Low |
| **Ease of use** | High/Medium | Medium | Medium/Low | Low |

\* High if exact match or conspecific in database, medium to low if not
\*\* If already trained it is extremely fast, but training can require high computational power
\*\*\* If enough signal in the sequence (See Janssen *et al.* 2018)

**Figure 2.**
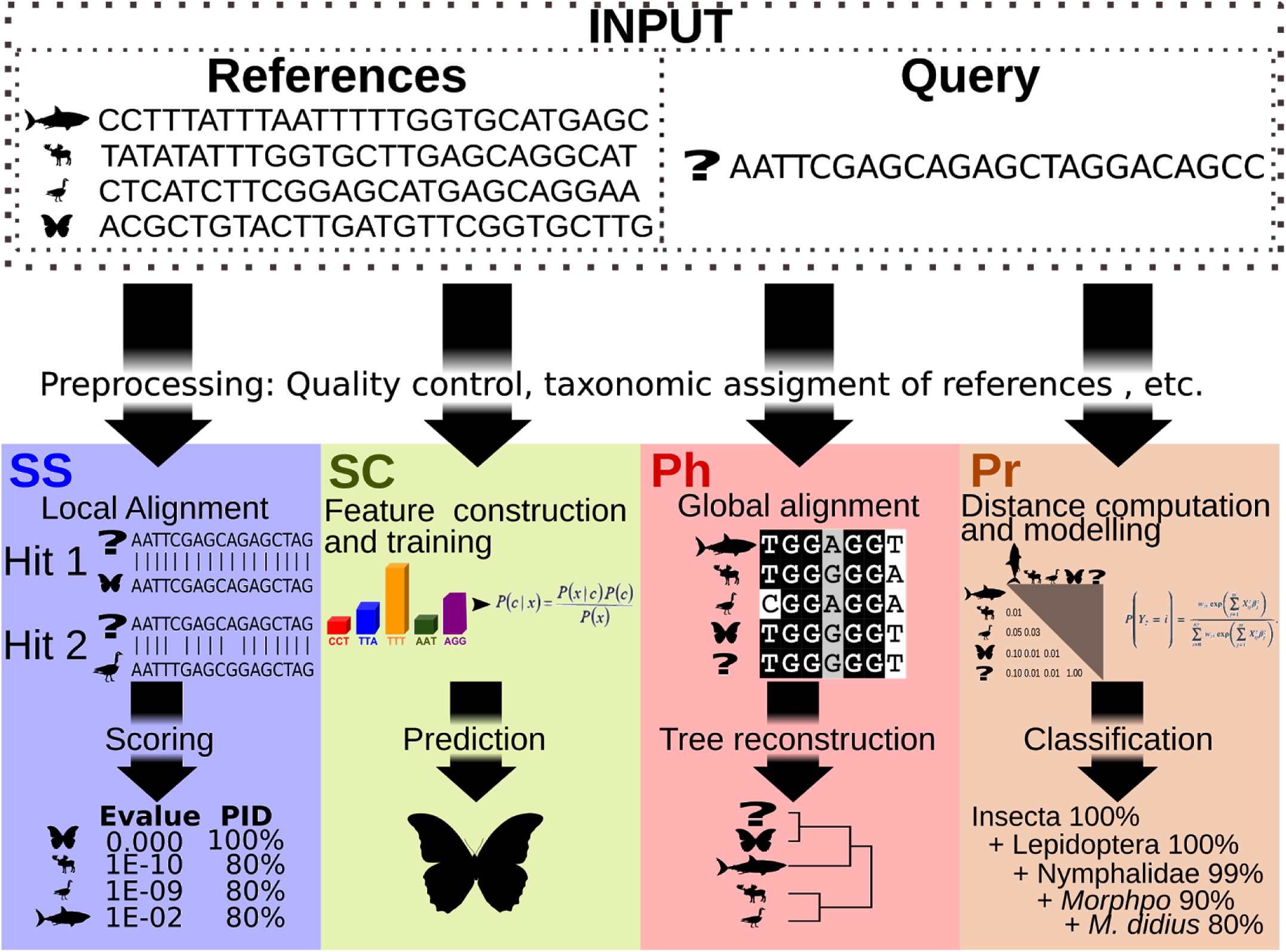
Overview of methodological approaches. Sequence similarity (SS) methods use local alignments to search for similarity between each query and the reference sequences. Sequence composition (SC) methods are trained by computing a k-mer frequency profile for each reference sequence, and then matching each query to this profile. Phylogenetic (PH) methods use global alignments including (or placing) the query in a phylogenetic tree. Probabilistic (PR) methods use a distance metric and then perform a hierarchical multinomial regression to estimate the certainty in the classification of each query at each taxonomic rank.

Despite this surge in method development, community adoption is lagging behind (Kumar & Dudley 2007). At least in part, this reflects the uncertainty in regard to which strategy will perform best in a particular situation. Our study addresses this conundrum by comprehensively assessing current taxonomic assignment tools including the optimization of parameters to maximize their accuracy. Accuracy of taxonomic assignments was evaluated using three types of *in vitro* mock communities with varying numbers of individuals and taxa representing compositional heterogeneity, defined as variability in community composition, both in number of species (between species variability) and in number of individuals (within species variability). This type of heterogeneity is important when assessing the quality of taxonomic assignment because such variability is inevitable in natural communities and it might confound predictions, and ignoring its effect can lead to false conclusions. In response to this concern, this study provides an unbiased benchmark of current tools for taxonomic assignment based on their performance with three mock communities with different degrees of heterogeneity. As such, it aims to provide guidance to the use of such methods, as well as landmarks for developers as they develop approaches to improve the accuracy of taxonomic assignments.

## METHODS

### Mock communities

Three types of mock communities were examined for this study: 1) Single individual per species, where multiple species are each represented by a single individual; 2) Multiple individuals per species, where multiple species are represented by a variable number of individuals (1- 23); and 3) Populations of a single species, consisting of multiple individuals of a single species. For the latter, populations of all species (Table S6) were pooled for community analyses.

Three taxonomically distinct mock communities were analyzed for the single individual per species. They included a zooplankton mock community consisting of 76 species representing four phyla (molluscs, rotifers, tunicates, crustaceans) belonging to 12 orders and 22 families (From Zhang *et al*. 2018b); a fish mock community with 41 species from 13 orders (This study, Supplementary Methods S1 and Supplementary Table S1, Dryad DOI wil be updated upon manuscript acceptance); and an insect mock community consisting of 365 species from 10 orders and 104 families (Braukmann *et al*. 2019).

The other two types of mock communities (multiple individuals per species, populations of a single species) were both assembled from the zooplankton set. The first was composed of multiple individuals (3 - 23) of 37 species (Supplementary Table S6). The populations of a single species community included four populations, each with several individuals of a single species (28 *Balanus crenatus*, 41 *Limnoperna fortunei*, 24 *Tortanus discaudatus*, 39 *Leptodora kindtii*; Supplementary Table S6).

This compositional heterogeneity allowed testing of the effectiveness of taxonomic assignment methods in systems with different levels of genetic variation, as it is typical for natural communities. Raw sequence data is available through Dryad (Zhang *et al*. 2018a). Our analysis focuses on two widely used COI metabarcoding markers: the Leray fragments (Leray *et al*. 2013) for the zooplankton and fish mock communities, and the MLepF1/LepR1 (Hebert *et al*. 2004) for the insect mock community.

### Quality control, merging, and denoising pipeline

To guarantee unbiased benchmarking, all mock communities were analyzed using identical steps: filtering, merging, and denoising. The quality of raw reads was assessed with the *quality plot* function implemented in the DADA2 pipeline (Callahan *et al*. 2016) but modified to automatically detect the low quality end trimming position. The automatic detection of trimming length was set to a mean quality score below 25. The resultant position was recorded and Cutadapt v1.18 (Martin 2011) was employed to trim each read along with adapters and primers using parameters shown in Supplementary Table S2.

To minimize variance introduced during preprocessing of reads, we employed the DADA2 (Callahan *et al*. 2016) pipeline for filtering and trimming, modelling sequencing error, dereplication, denoising, merging reads, amplicon sequence variant (ASV) table creation, and removal of chimeric sequences before input into the respective taxonomic assignment software (Supplementary Table S2). To exclude sequences generated by non-target priming, the length of predicted ASVs was restricted to +- 20bp of the expected fragment length: between 293 and 333bp for the Leray fragment (Leray *et al*. 2013), and between 387 and 427 for the insect fragment (Braukmann *et al*. 2019). ASVs composed of less than eight denoised sequences were removed to reduce noise and spurious detection and to match default parameters of programs such as UPARSE (Edgar 2013). The pipeline was written in R and is available at https://github.com/jshleap/TA_pipes/blob/master/dada2_pipe.R.

### Curating mock community composition

Although the original species composition and provenance of the DNA extract for each mock community was known, the actual species composition of the amplicon pool submitted for sequencing can differ from it. This difference can arise from PCR bias or from the loss of sequences during bioinformatic processing (Epp *et al*. 2012). Reads can also derive from unintended sources of DNA such as parasites, gut contents (Zhang *et al*. 2018b), or environmental DNA. To avoid such problems, we validated and corrected the composition of sequences recovered from each mock community to reflect its true composition through a combined screen employing a high stringency BLAST search (Altschul *et al*. 1997) and a phylogenetic approach. First, we created a local database with the known composition of each mock community by mining reference sequences from the NCBI database. For the fish mock community, we also supplemented the database with sequences of some expected species (Supplementary Methods S1). We then performed a nucleotide BLAST search of the mock communities, using the newly created database, restricting the search to an e-value of 0.001 and 90% identity. Hits with over 99.5% identity and 100% query coverage were deemed to derive from species in the assemblages and taxonomy was assigned directly from the BLAST hit. Sequences without assignment required further careful examination of homology and phylogenetic distance. For this, BLAST hits were sorted by percent identity and query coverage and the top three unique species hits were selected. A BLAST search for sequences of congeneric taxa was performed using the NCBI non-redundant nucleotide database. If available, up to three different congeneric species were retrieved. Sequences from the top three rows of the e-value-sorted BLAST table were retrieved and merged with both queries and congeneric sequence hits. Merged sequences were then aligned with MAFFT v7.310 (Katoh & Standley 2013) using the *auto* option. In downstream analysis, fully gapped sequences, and columns of sequences with over 90% gaps were removed. A maximum likelihood tree was constructed with FastTree (Price *et al*. 2010) using the GTR + gamma model, and all branches with less than 70% support in a Shimodaira-Hasegawa test were collapsed. Each query sequence clade was extracted ensuring that at most one species was retained (as per reference sequences) and an outlier test of branch lengths was performed. If no outlier was found, the clade was considered monophyletic for one species. If this species was part of the intended community, the assignment was automatically made. A second round of BLAST and tree reconstruction was performed for all clades and sequences that did not find a significant hit in the previous step, increasing the number of top hits to 30. Sequences that could not be reliably assigned manually to species level or that were identified as contaminants (e.g. fungi, synthetic constructs) were removed from the sequence array for the community. Supplementary Table S3 provides the resulting taxonomic diversity for each mock community. All steps were run using an in-house python script *process_mock.py* available at https://github.com/jshleap/TA_pipes/blob/master/process_mock.py with manual curation performed in an Ipython shell using the ete3 library (Huerta-Cepas *et al*. 2016).

### Assembly of reference databases

All methods for taxonomy assignment require comprehensive, well-annotated reference databases to perform effectively. COI sequences for this study were mined from NCBI GenBank, BOLD Systems (http://www.boldsystems.org; Ratnasingham & Hebert 2007), and MIDORi (http://reference-midori.info; Leray *et al*. 2018). Sequence duplicates were removed using SeqKit (Shen *et al*. 2016). Information on seven taxonomic levels (Kingdom, Phylum, Class, Order, Family, Genus, Species) was retrieved and added to the sequence header using TaxonKit (Shen & Xiong 2019). Only sequences with complete 7-level taxonomic information were retained. Because phylogenetic and probabilistic methods require a global alignment, only a subset of reference sequences was used. This subset was created by identifying the lowest taxonomic level shared by all curated sequences in each mock community and picking one representative per species. Each subsampled reference set was subsequently aligned using the Amplicons to Global Gene (A2G^2^) program which generates very large alignments while minimizing gaps (Hleap *et al*. 2020).

Subsampling strategies, like the one described above, are often used when the reference sequence database is large and phylogenetic taxon assignment is used (Chesters 2017; Hardge *et al*. 2018; Czech *et al*. 2019), thus providing a realistic use scenario.

### Inclusion of true negatives

In order to quantify cases of spurious detection (i.e. false detection of a species), we randomized (i.e. shuffled) sequences following Lindgreen et al. (2016). We randomly sampled 10% of the assembled, denoised, and chimera-free reads using Seqkit v0.12.0 (Shen *et al*. 2016), and shuffled these sequences keeping both mononucleotide composition (sampling from the average nucleotide frequency distribution) and di-residue composition (sampling for the distribution of nucleotide pairs), creating synthetic sequences with the same overall nucleotide composition as the original set. This process was performed with the esl-shuffle function of HMMER3 v.3.2.1 (Eddy 2011). These sequences were used to determine the number of true negatives in the query.

### Metrics for benchmarking

Four metrics were used to compare performance of the seven taxonomic assignment methods: true positives (TP), false positives (FP), false negatives (FN), and true negatives (TN). The TP are sequences known to derive from a species present in the mock community and assigned to this taxon. FP are sequences that were assigned to a species not present in the community, while FN are species known to be present but that had no sequence assigned to them. Finally, the TN are sequences present in the dataset that should not be assigned to any species. To compute TN, we determined how many of the randomized sequences were assigned to any of the 7 taxonomic levels and subtracted that value from the total randomized sequences in each mock community. We also measured five composite metrics:

The false discovery rate (FDR), the proportion of false predictions made, is calculated as:

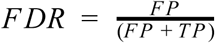

The true positive rate (TPR) or sensitivity is the proportion of true positive predictions for species present in the sample and is calculated as:

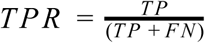

The positive predictive value (PPV) is the fraction of true positives from all predictions (label assignments) and is calculated as:

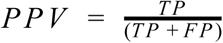

The F1-Score is the harmonic mean of TPR and PPV. It represents the overall measure of confidence in a prediction as a trade-off between TPR and PPV. It can be estimated by:

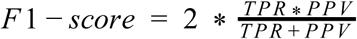

Finally, the Matthews correlation coefficient (MCC) corresponds to the correlation between the observed and predicted classification and is bounded by values ranging from −1 to 1. It is calculated as:

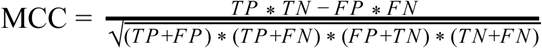

This metric is akin to the Pearson correlation coefficient between two variables and can be interpreted in a similar manner. If the Mathews correlation coefficient is 1, the two variables are perfectly correlated while a value of 0 means that the prediction is no better than random. Negative values indicate that the prediction is more frequently incorrect than expected by chance. A comparison of the metrics in other benchmarking efforts is provided in Supplementary Table S4, and a mock example of how each metric works can be found in the Supplementary Note 1.

Along with these classification metrics, the computational effort required to carry out each analysis was measured based on percentage of CPU utilization, the time taken to make a prediction and memory usage. The time measurements only considered the prediction step and not the resources required for training the machine learning approaches or constructing the reference databases for the sequence similarity methods.

### Strategy for selecting taxonomic assignment programs

For each taxonomic assignment category we evaluated two programs based on four criteria: 1) Frequency of use, as measured by the average number of citations per year over the last five years (Figure 1); 2) Type of assignment or category (as per Figure 2); 3) Novelty (i.e. programs without prior benchmarking); and 4) Applicability (i.e. programs with active support that are not marker specific). We employed two sequence similarity methods, Kraken2 (Wood & Salzberg 2014) and Basta (Kahlke & Ralph 2019); two sequence composition methods, QIIME q2-feature-classifier (Bokulich *et al*. 2018) and IDtaxa (Murali *et al*. 2018); and one probabilistic method, Protax (Somervuo *et al*. 2016; Axtner et al. 2019). The two most frequently cited phylogenetic methods are TIPP (Nguyen *et al*. 2014) and MLTreeMap (Stark *et al*. 2010), but neither have active development or support so were not evaluated. As a phylogenetic method, we chose HMMUFOTU (Zheng *et al*. 2018) because all other implementations (e.g. pplacer, EPA) in this category leave taxon assignments to postprocessing after phylogenetic placement. As a baseline reference and given that it is still widely used in metabarcoding studies, we also included BLAST top hits in the tests. Program descriptions and characteristics are provided in Supplementary Table S5.

### Optimization of parameters and evaluation of accuracy

Each program designated for evaluation has a set of adjustable parameters which can affect its performance in making taxonomic assignments. We optimized parameter selection for each program rather than adopting a one-size-fits-all approach. We did this by creating a linear search for each parameter employing the objective criterion of maximizing the Mathews correlation coefficient because it incorporates all basic metrics (see Metrics for benchmarking section). This selection of the best parameter set was done by performing a grid search with pseudo 3-fold cross validation, where one third of each mock community was selected at random to identify the best parameter values, while the final test was performed on the remaining two thirds of the community. With this procedure, some degree of overfitting can be accomodated. When multiple combinations of parameters yielded the same results, the least stringent parameter set was chosen to ensure generalizability of predictions. To test the effect of parameter optimization on accuracy of taxonomic assignment, we performed two multiple linear regressions with all the parameters as independent variables and the F1-score and Matthews correlation coefficient as response variables. With the estimated weights, we predicted the accuracy value for every combination of parameters. We then computed R^2^ between the real and predicted values as a proxy of fit for each one of the training folds (folds being each partition of the data) during cross validation. This R^2^ represents the proportion of the variance predictable from parameter usage, and therefore is a useful proxy for the effect of parameter tuning on accuracy.

## RESULTS

### Accuracy

We evaluated the Matthews correlation coefficient and F1-score (basic metrics and other compound metrics are provided in supplementary table S6) for each method at the family, genus, and species level as these are the key taxonomic ranks for ecological studies (Thiault *et al*. 2015; Wiese *et al*. 2016). QIIME2 generated the highest average Mathews correlation coefficient for accuracy of classification among the methods tested for each mock community, achieving average values of 0.99 (0.02 standard deviation), 0.89 (0.11 standard deviation), and 0.67 (0.09 standard deviation) for family, genus, and species, respectively (Figure 3 and Supplementary Table S6). The loss of accuracy towards the lower taxonomic ranks reflected an increase in the false discovery rate (increased false positives) rather than a lack of predictions (false negatives) (Supplementary Table S6). This pattern was true for all communities except the insect mock community where more false negatives were found but fewer false positives (Figure3 and Supplementary Table S6).

**Figure 3.**
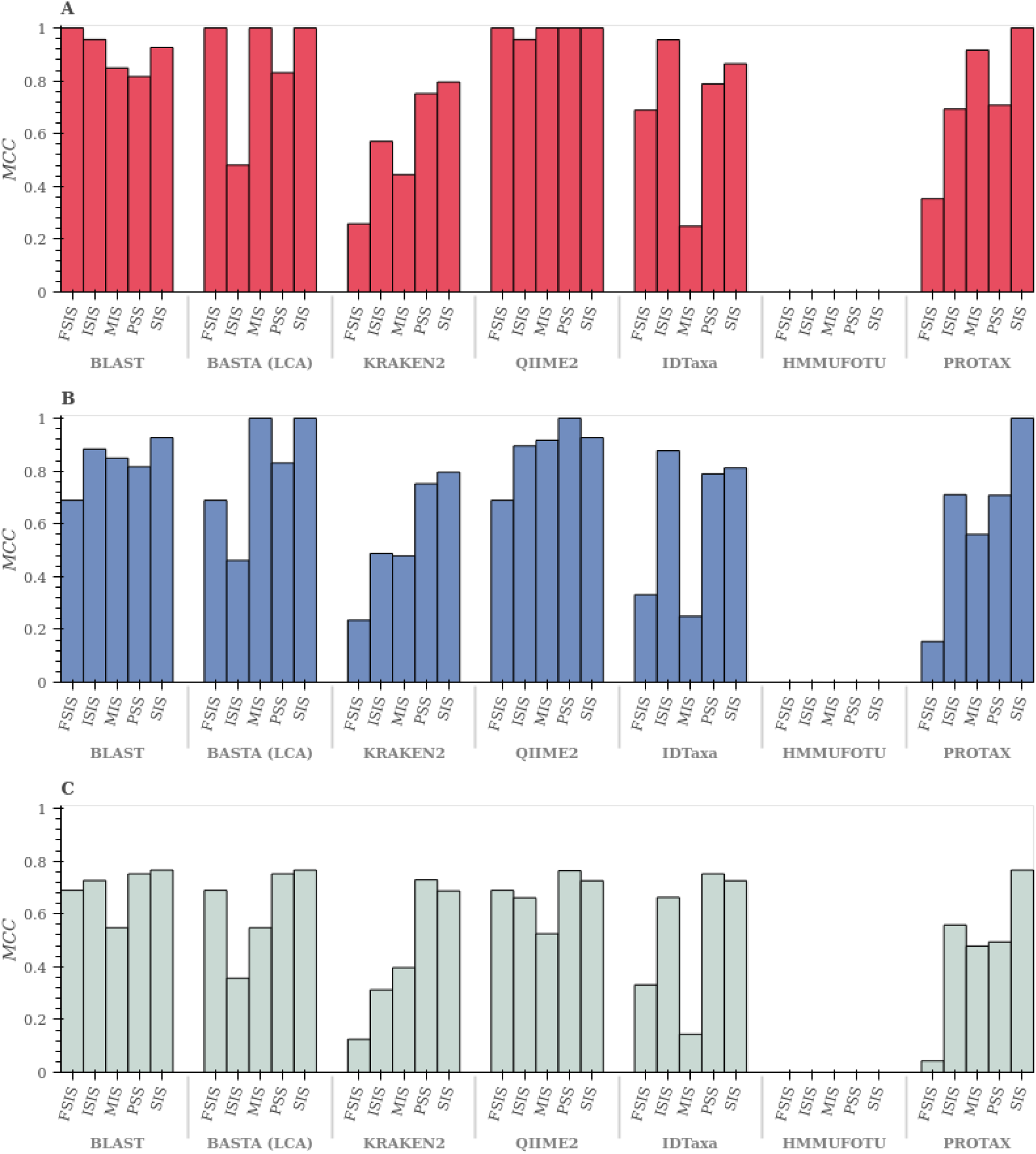
Matthews correlation coefficient (MCC) for each mock community for all assignment methods. Taxonomic assignment was examined at three levels: A) Family, B) Genus, and C) Species. FSIS: Fish single individual per species; ISIS: Insect single individual per species; MIS: Zooplankton multiple individuals per species; PSS: Zooplankton population of single species; SIS: Zooplankton single individual per species.

QIIME2 was closely followed by BLAST which had average Mathews correlation coefficient values of 0.91 (0.08 standard deviation - SD), 0.83 (0.09 SD), and 0.70 (0.09 SD) for family, genus, and species respectively (Figure 3 and Supplementary Table S6). BLAST did not produce false negatives for any mock community and no true negatives were predicted (Supplementary Table S6). Loss of accuracy at lower taxonomic levels again resulted from false positives, with the multiple individual per species community having the highest relative incidence of false positives (Supplementary Table S6).

In comparison to QIIME2 and BLAST, Kraken2 generated much lower average Mathews correlation coefficient values for family (MCC = 0.56, SD = 0.22), genus (MCC = 0.55, SD = 0.23), and species (MCC = 0.45, SD = 0.26) (Supplementary Table S6). Kraken did correctly avoid assigning taxonomy to the shuffled sequences, but produced many false negatives, especially for the fish community (e.g. it failed to assign *Catostomus commersonii* and *Notropis hudsoni* to the correct species), and some false positives particularly for the insect community (e.g. *Dioryctria abietella, Euura clitellata*; Supplementary Table S6).

The type of mock community did not have a substantial effect on the accuracy of either QIIME or BLAST as both methods had less than 0.1 standard deviation (SD) of the Mathews correlation coefficient (with the exception of QIIME in species with 0.12). All other methods showed high levels of variation in accuracy (> 0.2 SD except LCA which had a SD of 0.17 at the species-level) depending on the type of mock community. The zooplankton population of single species, and the single individual per species mock communities were the least variable and the fish mock community the most variable (Supplementary Table S6 and Supplementary figure S2). The probabilistic method (Pr) PROTAX, exhibited many false negatives especially for the fish community.

HMMUFOTU always assigned the shuffled sequences to a taxon, leading to no true negatives, and therefore rendering the Mathews correlation coefficient undefined (Figure 3). For reference, we provide the same analysis using the F1-score as an accuracy metric in the supplementary materials (Supplementary figure S2). As this metric does not consider true negatives, its performance is overestimated. Most methods showed similar trends in both the F1-score and the Mathews correlation coefficient analyses: QIIME and BLAST performed best while KRAKEN2 and HMMUFOTU were worst.

### Parameter optimization

Parameter optimization played an important role in accuracy of taxonomic assignments (Figure 4; supplementary tables S7-S13) for most methods. However, our results suggest that Kraken2 was the most affected (Figure 4) as evidenced by its higher overall R^2^ over multiple linear regressions with all the parameters as independent variables and the F1-score and Matthews correlation coefficient as response variables. Kraken2 showed an average R^2^ of 0.74 meaning that 74% of its variance is explained by parameter tuning. By contrast, the PROTAX (probabilistic method) confidence (conf) parameter had no influence on the predicted composition of any mock community regardless of parameter tuning (Figure 4 and supplementary table S11). A similar pattern was found for HMMUFOTU (phylogenetic method) with an average R^2^ of 0.05. However, there seemed to be a correlation with the type of mock community as the zooplankton single individual per species (for the Mathews correlation coefficient metric; Figure 4) responded best to optimization.

**Figure 4.**
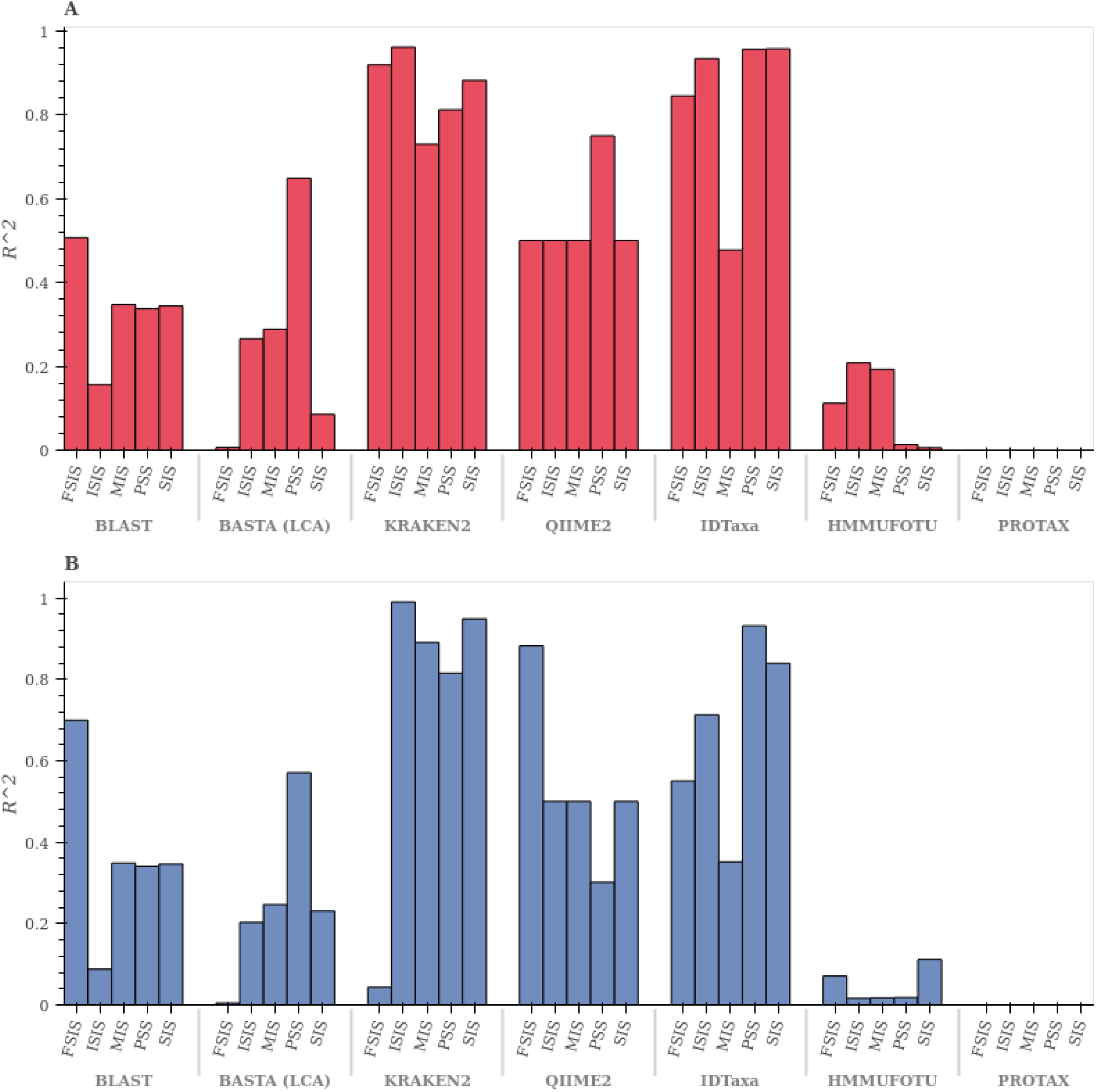
Distribution of R^2^ per method and mock community. Each R^2^ value is the fit between the predicted accuracy based on a multiple regression (accuracy as dependent variable and all the parameters as independent variables) and the actual accuracy obtained. A) F1-score; B) Mathews correlation coefficient (MCC).

The most parameterized method is BASTA’s last common ancestor-type software with five tunable parameters with 5 -11 levels for each (Supplementary Table S6). This requires an optimization run for more than 23,000 combinations of parameters. As a result, BASTA took the longest (>40 hours of compute time) to provide optimization results, despite being one of the fastest methods for single runs (Figure 5 and Supplementary Table S6). In terms of response to optimization, BASTA exhibited an average R^2^ of 0.25 (i.e. 25% of variance reflects parameter tuning; Tables S8a to S8e). It appears to be community dependent (Figure 4) as tuning had a more profound effect on the population of a single species community with an R^2^ of 0.57.

**Figure 5.**
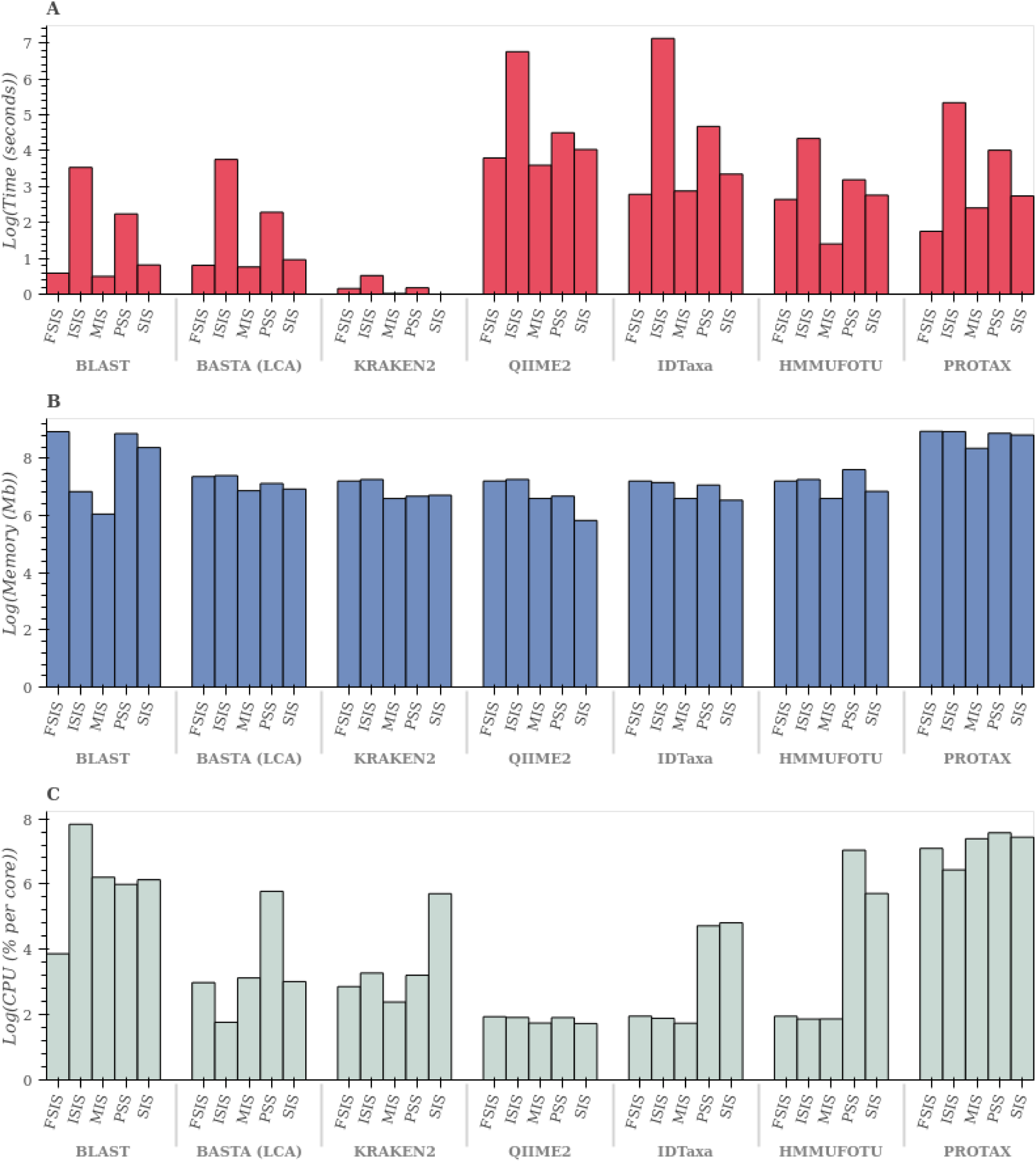
Overall performance of the programs evaluated for all mock communities. A) Time of execution in log(seconds); B) Memory used in log(Mb); C) CPU usage in log (percentage CPU usage per core). The y-axis is log transformed to aid visualization. FSIS: Fish single individual per species; ISIS: Insect single individual per species; MIS: Zooplankton multiple individuals per species; PSS: Zooplankton population of single species; SIS: Zooplankton single individual per species.

PROTAX, IDtaxa, and Kraken2 are the least parameterized methods with only one tunable parameter (Table 2), which translates into a smaller search space for the optimum value, and therefore potentially less time to determine the optimal solution. PROTAX however, did not respond to optimization as it showed similar Mathews correlation coefficient values for all levels of the tunable parameter (confidence threshold in this case; Table 2). The variance in accuracy for IDtaxa and Kraken2 could be explained by parameter optimization. Both programs might deliver improved accuracy when detailed parameter tuning is applied as they currently rely on just a single parameter.

**Table 2.** Parameter space for the optimization of the programs used.

| Program | Parameter | Space |
| --- | --- | --- |
| Blast (Top hit) | e-value | {1E-50, 1E-25, 1E-15, 1E-7, 1E-4, 1E-2} |
|  | Percent identity | {70, 75, 80, 85, 90, 95, 100} |
|  | Max target sequences | {1, 25, 50, 75, 100, 500} |
| BASTA (LCA) | e-value | {1E-50, 1E-25, 1E-15, 1E-7, 1E-4, 1E-2} |
|  | Percent identity | {70, 75, 80, 85, 90, 95, 100} |
|  | Minimum number of hits | {1, 2, 3, 4, 5, 6, 7, 8, 9, 10} |
|  | Maximum number of hits | {0, 1, 2, 3, 4, 5, 6, 7, 8, 9, 10} |
|  | Percentage of hits to use | {60, 70, 80, 90, 99} |
| Kraken2 | Confidence threshold | {0, 0.1, 0.2, 0.3, 0.4, 0.5, 0.6, 0.7, 0.8, 0.9, 1.0} |
| QIIME | Confidence threshold | {0.5, 0.6, 0.7, 0.8, 0.9, 1.0} |
| IDTAXA | Confidence threshold | {0.5, 0.6, 0.7, 0.8, 0.9, 1.0} |
|  | Bootstraps | {50, 100, 200, 400} |
|  | Minimum fraction of bootstraps | {0.9, 0.92, 0.94, 0.96, 0.98, 1.0} |
| HMMUFOTU | Seed | {1, 25, 50, 75, 100} |
|  | Max observed distance | {0, 0.1, 1, 10, 'inf'} |
|  | Max placement error | {1, 10, 20, 30, 40} |
|  | Branch length estimating method | {'unweighted', 'weighted'} |
|  | Method for calculating prior probability of a placement | {'uniform', 'height'} |
| PROTAX | Confidence threshold | {0.1, 0.05, 0.01, 0.001} |

### Overall performance

Kraken2 was the most time-efficient method, being several orders of magnitude faster than the other methods evaluated, reaching results in 0.5 sec on average (Figure 5A and Supplementary Table S6). BLAST and BASTA were slightly higher, generating results in 5.8 and 7.3 seconds averaged across the three mock communities. By contrast, IDtaxa and QIIME were almost three orders of magnitude slower than Kraken2, with 129.4 and 99.6 seconds on average across mock communities, respectively (Figure 5A and Supplementary Table S6).

In terms of memory, all programs had roughly similar requirements, but PROTAX and BLAST were the most memory intensive using an average of 6648 and 4054 megabytes of memory, respectively (Figure 5B and Supplementary Table S6) while QIIME required the least (924.3 Mb; Figure 5B and Supplementary Table S6) during the classification phase.

Another performance metric was CPU usage, the proportion of a single core used to execute the program. As most computers now contain multiple CPU cores, the percentage of usage can be higher than 100%. CPU usage differed substantially among mock communities, largely reflecting their varying number of sequences (Figure 5C). PROTAX and BLAST showed consistently high CPU usage, with 1414.8 and 784.3 average CPU percentage, respectively (Figure 5C and Supplementary Table S). On average, QIIME was the most CPU efficient software, using only 6% CPU on average.

## DISCUSSION

In this study we comprehensively assessed the performance of current taxonomic assignment software. It is often assumed that more sophisticated methods outperform BLAST in performance and accuracy of taxonomic assignments in almost every setting, adding to reports of false positives (Virgilio *et al*. 2010; Porter & Hajibabaei 2018). However, our results suggest that added complexity does not always yield significantly better results, especially if a mock community resembling the expected sample can be used for parameter adjustment.

### Classification Accuracy

Naive Bayes classifiers have been shown to outperform BLAST in certain settings (Rosen *et al*. 2011) but we only found this to be true for QIIME, suggesting that other classifiers are extremely sensitive to the reference database available, especially with respect to composition and the number of reference sequences per species (Schenekar *et al*. 2020). Although QIIME slightly outperformed BLAST, sequence similarity (SS) methods seemed more robust for highly heterogeneous and large databases. Both BASTA and Kraken2 use the last taxonomic common ancestor (LCA) strategy to assign taxonomy, and their performance might be underestimated in this study, since we tested each taxonomic level strictly. This means that even if these methods correctly identified a higher taxonomic level, it would be reported as a misclassification for the ranks evaluated. This reflects the heterogeneity in LCA predictions (i.e. not all taxonomic levels are classified to the same depth). If a study does not require strict taxonomic levels (i.e. species level prediction is not required in all assignments) then LCA methods might deliver higher accuracies than reported here. In fact, a LCA approach can be very appropriate when some taxa are undersampled, e.g. in exploratory analyses of diversity or environmental DNA studies. In such cases, the sensitivity of LCA approaches might reveal the presence of undersampled or rare species (e.g tropics; Bacci *et al*. 2018) at higher taxonomic levels (e.g. family) when a genus or species identification is not possible In our analysis, PROTAX exhibited low accuracy when making assignments for the fish mock community. This result might reflect the presence of records from hybrids or from misidentified specimens in the reference database as these would confound the regression classifier. It is certain that sequences derived from hybrids are present in reference databases such as the NCBI, and their annotation is extremely variable, making them likely to be present in most reference databases that have not been manually curated. Also, if the parental species of hybrids are used to identify species, the database might contain biases even after curation (Machida *et al*. 2017) and so will the classifier. This is an important factor to consider when dealing with taxonomic groups with high levels of hybridization such as fish. Taxonomic assignment in these cases should be inspected carefully and only highly curated databases should be used. However, introgression will affect the accuracy to the species level by increasing false positives, but might not affect other taxonomic ranks since introgression is rare above a genus level. Our results also revealed a high level of false negatives, likely reflecting the lack of enough reference sequences in each taxon. PROTAX requires at least two sequences of each taxon to model probabilities. Some of our reference databases only had one representative of certain species. This should not affect the genus or family level, but we observed instances where the genus and family were not assigned, although the reference database included multiple entries for congeners and confamilials.

PROTAX does have the advantage of informing the classifier of the likelihood of finding a given group (Axtner et al. 2019), e.g. if a species is present in the region under study. We did not test this feature since this information is rarely available for metabarcoding or ecological studies in understudied areas (e.g,. tropics) or taxonomic groups. We also deemed it not appropriate for a mock community study in order to avoid “observer” bias in already biased reference sets (Troudet *et al*. 2017). However, when prior information concerning expected diversity is available, PROTAX might have a better performance than achieved in our study.

Our results indicate that sample composition strongly affects the capacity of classifiers to make correct taxonomic assignments, an observation also reported in microbial metabarcoding (Yeoh *et al*. 2019). At the species level, accuracy across all methods for the multiple individuals per species mock community was low, perhaps due to confounding stemming from its higher genetic diversity per species (Supplementary Figure S2). This is important as most metabarcoding bulk samples are taxonomically heterogeneous and genetically diverse (Evans *et al*. 2017).

A high number of hybrids in databases, the relative low taxonomic diversity in public repositories, and a dynamic taxonomy (Mora *et al*. 2008; Vavalidis *et al*. 2019) perturb classification, and create high uncertainty of results (Supplementary Figure S2). This exemplifies the main sources of error in taxonomic assignment: mis-annotations in reference databases and the incomplete representation of taxa in them (Troudet *et al*. 2017; Leray *et al*. 2018; Macheriotou *et al*. 2019).

### Classification parameter optimization

Although it is intuitive that parameter optimization should improve the accuracy of predictions (Bokulich *et al*. 2018), many studies employ previously published or default parameters (e.g. Anantharaman *et al*. 2016; Rodríguez-Martínez *et al*. 2020), thereby potentially propagating errors. Our data show that the accuracy of most methods is strongly influenced by parameter optimization, particularly in Kraken2, IDtaxa, and BASTA. The assembly of a mock community closely resembling the expected sample followed by parameter tuning before making taxonomic assignments could yield a more accurate description of the community by reducing false positives and false negatives (Zhang *et al*. 2018b; Braukmann *et al*. 2019).

PROTAX was the sole method that showed no response to parameter tuning. This lack of relationship might be explained by the robust parametrization of the multinomial model, where the probability values lie outside the bounds of the tested thresholds. This strength becomes very important when no ground truthing is available for samples.

### Classification performance

During the classification of the mock communities, all methods completed analysis rapidly (39 ± 112 seconds). Kraken2 was most rapid reflecting its optimization for data heavy metagenomic studies (Wood *et al*. 2019). BLAST-based methods (BLAST top hit and BASTA LCA) were more efficient than the SC methods, but BLAST scales poorly (Porter & Hajibabaei 2018). The three methods based on sequence composition (Idtaxa, QIIME) are least scalable, followed by PROTAX. This implies that parameter optimization should be restricted to a smaller set, which comes with the caveat that the optimum might not be found in a realistic timeframe.

Run times gain importance as reference databases and query sets grow, especially with parameter optimization. Some methods lack an off-the-shelf way to parallelize a run. For example, the LCA implementation (BASTA; Kahlke & Ralph 2019) involves a database that cannot be concurrently accessed, making BASTA single threaded (one query at a time). Despite this limitation, it ran rapidly, but parameter optimization is costly because of the lack of parallelization and number of parameters that can be tuned.

All programs had similar memory requirements (Figure 5B) except BLAST and PROTAX, which both required one more order of magnitude of memory during classification. PROTAX memory consumption remained high regardless of the size of the mock community or the number of reference sequences. Memory use in BLAST varied strongly with both factors, especially the number of reference sequences. A similar pattern was observed for CPU usage as PROTAX and BLAST required more than the other methods (between one and four orders of magnitude). This metric is important when multiple instances of programs are being run on a CPU, as this affects the run time of all active software. QIIME seems to be CPU-load independent which suggests that multiple instances of it can be run on the same CPU with little impact on individual run times.

### Training stage

So far we have only considered performance metrics at the classification stage. However, the performance during the training phase can also be a limiting factor. Sequence similarity programs are very fast during database creation and can process several million sequences (i.e. NCBI nt database) within a few minutes. By comparison, sequence composition programs are computationally expensive, with IDtaxa being the most extreme in this regard. During this study, QIIME required more than 500 Gb of RAM for 2+ days for our training set, while IDtaxa needed similar memory for 3+ days. These constraints can be overcome by using pre-trained references (e.g SILVA, GREENGENES), or smaller custom references. This comes at the potential cost of not discovering certain taxa and creating more false positives from a prediction of a confamilial instead of the target species. A possible strategy to deal with large reference databases is to first run BLAST/LCA methods on a higher taxonomic rank, assign the references to the identified families, and then create family-specific reference databases. However, even this approach can fail in extremely diverse families, and further subsets might be required.

Both phylogenetic and probabilistic (at least PROTAX) software also use extensive computational resources at the training phase. In addition, both require a global sequence alignment, and most alignment programs cannot deal with more than a few thousand sequences (Sievers et al., 2011). Alternative strategies to improve input alignment quality include anchoring the alignment to a particular region of the gene (e.g. COI) and to the amplicon within that gene (Hleap *et al*. 2020). What remains for both PROTAX and HMMUFOTU is an extensive training/tree building step after alignment that can be time and memory consuming. For HMMUFOTU, the computational impact is lower because of efficient phylogenetic software, such as FastTree (Price *et al*. 2010). However, these methods are less accurate in determining phylogenetic relationships and can therefore compromise downstream taxonomic assignments (Zhou *et al*. 2018). It is possible that the uncertainty introduced during the estimation of a very large phylogenetic tree with a fast heuristic such as the one used in FastTree is one reason why HMMUFOTU performed poorly in our study.

Following Gardner et al (2019), we have reported the most used accuracy metrics to aid future benchmarking, and made a neutral comparison (Boulesteix *et al*. 2013). Furthermore, of other neutral benchmarking studies (Bazinet & Cummings 2012; Lindgreen *et al*. 2016; Siegwald *et al*. 2017; Almeida *et al*. 2018; Gardner *et al*. 2019), ours is the only one to include manually curated real mock communities, and the addition of true negatives through sequence shuffling. To our knowledge, this is also the only neutral benchmark for amplicon sequencing in eukaryotes and it includes the largest real mock community to date, providing insights into the effect that query and reference sequence heterogeneity has on taxonomic assignments.

## CONCLUSIONS

Despite intensive research, and incremental improvements in software, the taxonomic assignment of sequence data remains challenging. As global biodiversity is under threat, it is important to gain the capacity to accurately determine alpha diversity. We show that for higher taxonomic ranks (e.g. family) current methods have roughly similar capacity to generate accurate predictions, but there is significant room for improvement at the genus and species levels. Achieving “exact” assignments is impossible at the species level because taxonomy is volatile, and because species are dynamic entities (both conceptually and genetically). However, there are a series of actions that help to minimize mis-assignments at all levels. Firstly, increased parameterization and strengthened curation of reference databases is of paramount importance for all identification software. Secondly, the construction of a mock community that corresponds to the diversity of the system under study has an important impact on parameter choice and overall taxonomic assignment accuracy. Thirdly, QIIME, BLAST or LCA methods, in conjunction with aforementioned parameter tuning, currently seem the best approaches for generating accurate taxonomic assignments.

## ACKNOWLEDGEMENTS

We thank Panu Somervuo for his help with the script for PROTAX and members of the Cristescu lab for fruitful discussions. We also thank Charles Xu and Jitka Krejci for input on the work and comments on this manuscript. We are also grateful to L. Veilleux, N. Vachon, H. Massé, and N. Tessier from the Ministère des Forêts de la Faune et des Parcs (Québec) for providing fish tissue samples. Compute Canada provided the HPC resources to train some software employed in this study. This research was supported by the Food from Thought: Agricultural Systems for a Healthy Planet Initiative, funded by the Canada First Research Excellence Fund (CFREF).

## REFERENCES

Almeida A, Mitchell AL, Tarkowska A, Finn RD (2018) Benchmarking taxonomic assignments based on 16S rRNA gene profiling of the microbiota from commonly sampled environments. GigaScience, 7. DOI: 10.1093/gigascience/giy054

Altschul SF, Madden TL, Schäffer AA et al. (1997) Gapped BLAST and PSI-BLAST: a new generation of protein database search programs. Nucleic Acids Research, 25, 3389–3402. DOI: 10.1093/nar/25.17.3389

Anantharaman K, Breier JA, Dick GJ (2016) Metagenomic resolution of microbial functions in deep-sea hydrothermal plumes across the Eastern Lau Spreading Center. The ISME Journal, 10, 225–239. DOI: 10.1038/ismej.2015.81

Axtner J, Crampton-Platt A, Hoerig LA et al. (2019) An efficient and robust laboratory workflow and tetrapod database for larger scale environmental DNA studies. GigaScience, 8(4). DOI: 10.1093/gigascience/giz029

Bacci LF, Amorim AM, Michelangeli FA, Goldenberg R (2018) Increased sampling in under-collected areas sheds new light on the diversity and distribution of *Bertolonia,* an Atlantic forest endemic genus. Systematic Botany, 43, 767–792. DOI: 10.1600/036364418X697490.

Barbera P, Kozlov AM, Czech L et al. (2019) EPA-ng: Massively Parallel Evolutionary Placement of Genetic Sequences. Systematic Biology, 68, 365–369. DOI: 10.1093/sysbio/syy054.

Bazinet AL, Cummings MP (2012) A comparative evaluation of sequence classification programs. BMC Bioinformatics, 13, 92. DOI: 10.1186/1471-2105-13-92.

Bokulich NA, Kaehler BD, Rideout JR et al. (2018) Optimizing taxonomic classification of marker-gene amplicon sequences with QIIME 2’s q2-feature-classifier plugin. Microbiome, 6, 90. doi: https://doi.org/10.1186/s40168-018-0470-z.

Boulesteix A-L, Lauer S, Eugster MJA (2013) A plea for neutral comparison studies in computational sciences. Plos One, 8, e61562. DOI: 10.1371/journal.pone.0061562.

Braukmann TWA, Ivanova NV, Prosser SWJ et al. (2019) Metabarcoding a diverse arthropod mock community. Molecular ecology resources, 19, 711–727. DOI: 10.1111/1755-0998.13008.

Callahan BJ, McMurdie PJ, Rosen MJ et al. (2016) DADA2: High-resolution sample inference from Illumina amplicon data. Nature Methods, 13, 581–583. DOI: 10.1038/nmeth.3869.

Chesters D (2017) Construction of a species-level tree of life for the insects and utility in taxonomic profiling. Systematic Biology, 66, 426–439. DOI: 10.1093/sysbio/syw099.

Clemente JC, Jansson J, Valiente G (2011) Flexible taxonomic assignment of ambiguous sequencing reads. BMC Bioinformatics, 12, 8. DOI: 10.1186/1471-2105-12-8

Czech L, Barbera P, Stamatakis A (2019) Methods for automatic reference trees and multilevel phylogenetic placement. Bioinformatics, 35, 1151–1158. DOI: 10.1093/bioinformatics/bty767.

Eddy SR (2011) Accelerated profile HMM searches. PLoS Computational Biology, 7, e1002195. DOI: 10.1371/journal.pcbi.1002195.

Edgar RC (2013) UPARSE: highly accurate OTU sequences from microbial amplicon reads. Nature Methods, 10, 996–998.

Epp LS, Boessenkool S, Bellemain EP et al. (2012) New environmental metabarcodes for analysing soil DNA: potential for studying past and present ecosystems. Molecular Ecology, 21, 1821–1833. DOI: 10.1038/nmeth.2604.

Evans NT, Li Y, Renshaw MA et al. (2017) Fish community assessment with eDNA metabarcoding: effects of sampling design and bioinformatic filtering. Canadian Journal of Fisheries and Aquatic Sciences, 74, 1362–1374. DOI: 10.1139/cjfas-2016-0306.

Folmer O, Black M, Hoeh W, Lutz R, Vrijenhoek R (1994) DNA primers for amplification of mitochondrial cytochrome c oxidase subunit I from diverse metazoan invertebrates. Molecular Marine Biology and Biotechnology, 3, 294–299.

Fosso B, Pesole G, Rosselló F, Valiente G (2018) Unbiased taxonomic annotation of metagenomic samples. Journal of Computational Biology, 25, 348–360. DOI: 10.1089/cmb.2017.0144.

Gardner PP, Watson RJ, Morgan XC et al. (2019) Identifying accurate metagenome and amplicon software via a meta-analysis of sequence to taxonomy benchmarking studies. PeerJ, 7, e6160. DOI: 10.7717/peerj.6160

Gillet B, Cottet M, Destanque T et al. (2018) Direct fishing and eDNA metabarcoding for biomonitoring during a 3-year survey significantly improves number of fish detected around a South East Asian reservoir. Plos One, 13, e0208592. DOI: 10.1371/journal.pone.0208592.

Guo Z, Zhang L, Li Y (2010) Increased dependence of humans on ecosystem services and biodiversity. Plos One, 5(10), e13113. DOI: 10.1371/journal.pone.0013113.

Hardge K, Neuhaus S, Kilias ES et al. (2018) Impact of sequence processing and taxonomic classification approaches on eukaryotic community structure from environmental samples with emphasis on diatoms. Molecular Ecology Resources, 18, 204–216. DOI: 10.1111/1755-0998.12726.

Hebert PDN, Penton EH, Burns JM, Janzen DH, Hallwachs W (2004) Ten species in one: DNA barcoding reveals cryptic species in the neotropical skipper butterfly *Astraptes fulgerator*. Proceedings of the National Academy of Sciences of the United States of America, 101, 14812–14817. DOI: 10.1073/pnas.0406166101.

Heeger F, Bourne EC, Baschien C et al. (2018) Long-read DNA metabarcoding of ribosomal RNA in the analysis of fungi from aquatic environments. Molecular Ecology Resources, 18, 1500–1514. DOI: 10.1111/1755-0998.12937.

Hernández-Triana LM, Prosser SW, Rodríguez-Perez MA et al. (2014) Recovery of DNA barcodes from blackfly museum specimens (Diptera: Simuliidae) using primer sets that target a variety of sequence lengths. Molecular Ecology Resources, 14, 508–518. DOI: 10.1111/1755-0998.12208.

Hleap JS, Cristescu ME, Steinke D (2020) A2G ^2^ : A Python wrapper to perform very large alignments in semi-conserved regions. BioRxiv .2020.05.21.109009; DOI: 10.1101/2020.05.21.109009.

Huerta-Cepas J, Serra F, Bork P (2016) ETE 3: reconstruction, analysis, and visualization of phylogenomic data. Molecular Biology and Evolution, 33, 1635–1638. DOI: 10.1093/molbev/msw046.

Ivanova NV, Dewaard JR, Hebert PDN (2006) An inexpensive, automation-friendly protocol for recovering high-quality DNA. Molecular Ecology Notes, 6, 998–1002. DOI: 10.1111/j.1471-8286.2006.01428.x.

Janssen S, McDonald D, Gonzalez A et al. (2018) Phylogenetic placement of exact amplicon sequences improves associations with clinical information. mSystems, 3(3), e00021–18. DOI: 10.1128/mSystems.00021-18.

Kahlke T, Ralph PJ (2019) BASTA – Taxonomic classification of sequences and sequence bins using last common ancestor estimations. Methods in Ecology and Evolution, 10, 100–103. DOI: 10.1111/2041-210X.13095.

Katoh K, Standley DM (2013) MAFFT multiple sequence alignment software version 7: improvements in performance and usability. Molecular Biology and Evolution, 30, 772–780. DOI: 10.1093/molbev/mst010.

Lan Y, Wang Q, Cole JR, Rosen GL (2012) Using the RDP classifier to predict taxonomic novelty and reduce the search space for finding novel organisms. Plos One, 7, e32491. DOI: 10.1371/journal.pone.0032491.

Leray M, Ho S-L, Lin I-J, Machida RJ (2018) MIDORI server: a webserver for taxonomic assignment of unknown metazoan mitochondrial-encoded sequences using a curated database. Bioinformatics, 34, 3753–3754. DOI: 10.1093/bioinformatics/bty454.

Leray M, Yang JY, Meyer CP et al. (2013) A new versatile primer set targeting a short fragment of the mitochondrial COI region for metabarcoding metazoan diversity: application for characterizing coral reef fish gut contents. Frontiers in Zoology, 10, 34. DOI: 10.1186/1742-9994-10-34.

Lindgreen S, Adair KL, Gardner PP (2016) An evaluation of the accuracy and speed of metagenome analysis tools. Scientific Reports, 6, 19233. DOI: 10.1038/srep19233.

Macheriotou L, Guilini K, Bezerra TN et al. (2019) Metabarcoding free-living marine nematodes using curated 18S and CO1 reference sequence databases for species-level taxonomic assignments. Ecology and Evolution, 9, 1211–1226. DOI: 10.1002/ece3.4814.

Machida RJ, Leray M, Ho S-L, Knowlton N (2017) Metazoan mitochondrial gene sequence reference datasets for taxonomic assignment of environmental samples. Scientific Data, 4, 170027. doi: https://doi.org/10.1038/sdata.2017.27.

Martin M (2011) Cutadapt removes adapter sequences from high-throughput sequencing reads. EMBnet.journal, 17, 10. DOI: 10.14806/ej.17.1.200

Mitra S, Stärk M, Huson DH (2011) Analysis of 16S rRNA environmental sequences using MEGAN. BMC Genomics, 12 Suppl 3, S17. doi: https://doi.org/10.1186/1471-2164-12-S3-S17.

Mizrahi-Man O, Davenport ER, Gilad Y (2013) Taxonomic classification of bacterial 16S rRNA genes using short sequencing reads: evaluation of effective study designs. Plos One, 8, e53608. DOI: 10.1371/journal.pone.0053608.

Mora C, Tittensor DP, Myers RA (2008) The completeness of taxonomic inventories for describing the global diversity and distribution of marine fishes. Proceedings. Biological Sciences, 275, 149–155. DOI: 10.1098/rspb.2007.1315.

Munch K, Boomsma W, Huelsenbeck JP, Willerslev E, Nielsen R (2008) Statistical assignment of DNA sequences using Bayesian phylogenetics. Systematic Biology, 57, 750–757. DOI: 10.1080/10635150802422316.

Murali A, Bhargava A, Wright ES (2018) IDTAXA: a novel approach for accurate taxonomic classification of microbiome sequences. Microbiome, 6, 140. DOI: 10.1186/s40168-018-0521-5.

Nguyen N-P, Mirarab S, Liu B, Pop M, Warnow T (2014) TIPP: taxonomic identification and phylogenetic profiling. Bioinformatics, 30, 3548–3555. DOI: 10.1093/bioinformatics/btu721.

Porter TM, Hajibabaei M (2018) Scaling up: A guide to high-throughput genomic approaches for biodiversity analysis. Molecular Ecology, 27, 313–338. DOI: 10.1111/mec.14478.

Price MN, Dehal PS, Arkin AP (2010) FastTree 2 — approximately maximum-likelihood trees for large alignments. Plos One, 5, e9490. DOI: 10.1371/journal.pone.0009490.

Ratnasingham S, Hebert PDN (2007) BOLD: The Barcode of Life Data System (http://www.barcodinglife.org). Molecular Ecology Notes, 7, 355–364. DOI: 10.1111/j.1471-8286.2007.01678.x.

Richardson RT, Bengtsson-Palme J, Johnson RM (2017) Evaluating and optimizing the performance of software commonly used for the taxonomic classification of DNA metabarcoding sequence data. Molecular Ecology Resources, 17, 760–769. DOI: 10.1111/1755-0998.12628.

Rodríguez-Martínez R, Leonard G, Milner DS et al. (2020) Controlled sampling of ribosomally active protistan diversity in sediment-surface layers identifies putative players in the marine carbon sink. The ISME Journal. 14(4), 984–998. DOI: 10.1038/s41396-019-0581-y.

Rosen GL, Reichenberger ER, Rosenfeld AM (2011) NBC: the Naive Bayes Classification tool webserver for taxonomic classification of metagenomic reads. Bioinformatics, 27, 127–129. DOI: 10.1093/bioinformatics/btq619.

Schenekar T, Schletterer M, Lecaudey LA, Weiss SJ (2020) Reference databases, primer choice, and assay sensitivity for environmental metabarcoding: Lessons learnt from a re-evaluation of an eDNA fish assessment in the Volga headwaters. River Research and Applications. 2020, 1–10. DOI: 10.1002/rra.3610f

Shen W, Le S, Li Y, Hu F (2016) SeqKit: A cross-platform and ultrafast toolkit for FASTA/Q File manipulation. Plos One, 11, e0163962. DOI: 10.1371/journal.pone.0163962.

Shen W, Xiong J (2019) TaxonKit: a cross-platform and efficient NCBI taxonomy toolkit. BioRxiv, 513523. doi: https://doi.org/10.1101/513523.

Shokralla S, Porter TM, Gibson JF et al. (2015) Massively parallel multiplex DNA sequencing for specimen identification using an Illumina MiSeq platform. Scientific Reports, 5, 9687. DOI: 10.1038/srep09687.

Siegwald L, Touzet H, Lemoine Y et al. (2017) Assessment of common and emerging bioinformatics pipelines for targeted metagenomics. Plos One, 12, e0169563. DOI: 10.1371/journal.pone.0169563.

Somervuo P, Koskela S, Pennanen J, Henrik Nilsson R, Ovaskainen O (2016) Unbiased probabilistic taxonomic classification for DNA barcoding. Bioinformatics, 32, 2920–2927. DOI: 10.1093/bioinformatics/btw346.

Stark M, Berger SA, Stamatakis A, von Mering C (2010) MLTreeMap--accurate Maximum Likelihood placement of environmental DNA sequences into taxonomic and functional reference phylogenies. BMC Genomics, 11, 461. DOI: 10.1186/1471-2164-11-461.

Thiault L, Bevilacqua S, Terlizzi A, Claudet J (2015) Taxonomic relatedness does not reflect coherent ecological response of fish to protection. Biological Conservation, 190, 98–106. DOI: 10.1016/j.biocon.2015.06.002.

Troudet J, Grandcolas P, Blin A, Vignes-Lebbe R, Legendre F (2017) Taxonomic bias in biodiversity data and societal preferences. Scientific Reports, 7, 9132. DOI: 10.1038/s41598-017-09084-6.

Vavalidis, Zogaris, Economou, Kallimanis, Bobori (2019) Changes in fish taxonomy affect freshwater biogeographical regionalisations: Insights from Greece. Water, 11, 1743. DOI: 10.3390/w11091743.

Virgilio M, Backeljau T, Nevado B, De Meyer M (2010) Comparative performances of DNA barcoding across insect orders. BMC Bioinformatics, 11, 206. DOI: 10.1186/1471-2105-11-206.

Wang Q, Garrity GM, Tiedje JM, Cole JR (2007) Naive Bayesian classifier for rapid assignment of rRNA sequences into the new bacterial taxonomy. Applied and Environmental Microbiology, 73, 5261–5267. DOI: 10.1128/AEM.00062-07.

Wiese R, Renaudie J, Lazarus DB (2016) Testing the accuracy of genus-level data to predict species diversity in Cenozoic marine diatoms. Geology, 44, 1051–1054. DOI: 10.1130/G38347.1.

Wood DE, Lu J, Langmead B (2019) Improved metagenomic analysis with Kraken 2. Genome Biology, 20, 257. DOI: 10.1186/s13059-019-1891-0.

Wood DE, Salzberg SL (2014) Kraken: ultrafast metagenomic sequence classification using exact alignments. Genome Biology, 15, R46. DOI: 10.1186/gb-2014-15-3-r46.

Yeoh YK, Chen Z, Hui M et al. (2019) Impact of inter- and intra-individual variation, sample storage and sampling fraction on human stool microbial community profiles. PeerJ, 7, e6172. DOI: 10.7717/peerj.6172.

Zhan A, Hulák M, Sylvester F et al. (2013) High sensitivity of 454 pyrosequencing for detection of rare species in aquatic communities. Methods in ecology and evolution / British Ecological Society, 4, 558–565. DOI: 10.1111/2041-210X.12037.

Zhang GK, Chain FJJ, Abbott CL, Cristescu ME (2018a) Data from: Metabarcoding using multiplexed markers increases species detection in complex zooplankton communities. Dryad Digital Repository. DOI: 10.5061/dryad.m83jc20

Zhang GK, Chain FJJ, Abbott CL, Cristescu ME (2018b) Metabarcoding using multiplexed markers increases species detection in complex zooplankton communities. Evolutionary Applications, 11, 1901–1914. DOI: 10.1111/eva.12694.

Zheng Q, Bartow-McKenney C, Meisel JS, Grice EA (2018) HmmUFOtu: An HMM and phylogenetic placement based ultra-fast taxonomic assignment and OTU picking tool for microbiome amplicon sequencing studies. Genome Biology, 19, 82. DOI: 10.1186/s13059-018-1450-0.

Zhou X, Shen X-X, Hittinger CT, Rokas A (2018) Evaluating Fast Maximum Likelihood-Based Phylogenetic Programs using empirical phylogenomic data sets. Molecular Biology and Evolution, 35, 486–503. DOI: 10.1093/molbev/msx302.

