## Supplementary Material for "Assessment of current taxonomic assignment strategies for metabarcoding eukaryotes"

### Supplementary Materials for the manuscript “Assessment of current taxonomic assignment strategies for metabarcoding eukaryotes: Insights from mock communities”

#### Table of Contents:

**Supplementary figures** 3

Figure S1. F1-score (F1S) in each of the mock communities for all methods evaluated. 3

Figure S2. Box plot of the MCC for each of the mock communities. 4

**Supplementary Tables** 5

Table S1. Fish Mock community composition. 5

Table S2. Parameters used in cutadapt and DADA2 pipeline 6

Table S3. Realized community in the mock after curation. 7

Table S4. Accuracy metrics used in benchmarking directly or indirectly taxonomic assignment software. 8

Table S5. Description of programs selected for the taxonomic assignment benchmark 8

Table S6. Performance and accuracy of the 6 programs evaluated. 9

Table S7. Optimization values for the parameters for Blast. 12

Table S8a. Optimization values for the parameters for LCA on the fish mock community (FSIS). 12

Table S8b. Optimization values for the parameters for LCA on the Insect mock community (ISIS). 12

Table S8c. Optimization values for the parameters for LCA on the zooplankton multiple individuals per species community (MIS). 12

Table S8d. Optimization values for the parameters for LCA on the zooplankton population single species (PSS) community. 12

Table S8e. Optimization values for the parameters for LCA on the zooplankton single individual per species mock community (SIS). 13

Table S9. Optimization values for the parameters for QIIME. 13

Table S10. Optimization values for the parameters for Kraken. 13

Table S11. Optimization values for the parameters for HMMUFOTU. 13

Table S12. Optimization values for the parameters for IDtaxa. 14

Table S13. Optimization values for the parameters for Protax. 14

**Supplementary methods** 14

**Methods S1** 15

Summary of methods 15

Sampling and identification of fish 15

Library preparation and next-generation sequencing (NGS) 16

Availability 16

**Supplementary notes** 17

Supplementary Note S1 17
On the complexity and accuracy gain of the methods evaluated 17
On parameter optimization 18
On the compositional heterogeneity 18
On the Accuracy metrics 19
On the practical context of the method’s performance 20
**References 20**

#### Supplementary figures

**Figure S1.** F1-score (F1S) in each of the mock communities for all methods evaluated.

A) Family level assignment; B) Genus level assignment; C) Species level assignment. FSIS: Fish single individual per species; ISIS: Insect single individual per species; MIS: Zooplankton multiple individuals per species; PSS: Zooplankton population of single species; SIS: Zooplankton single individual per species.

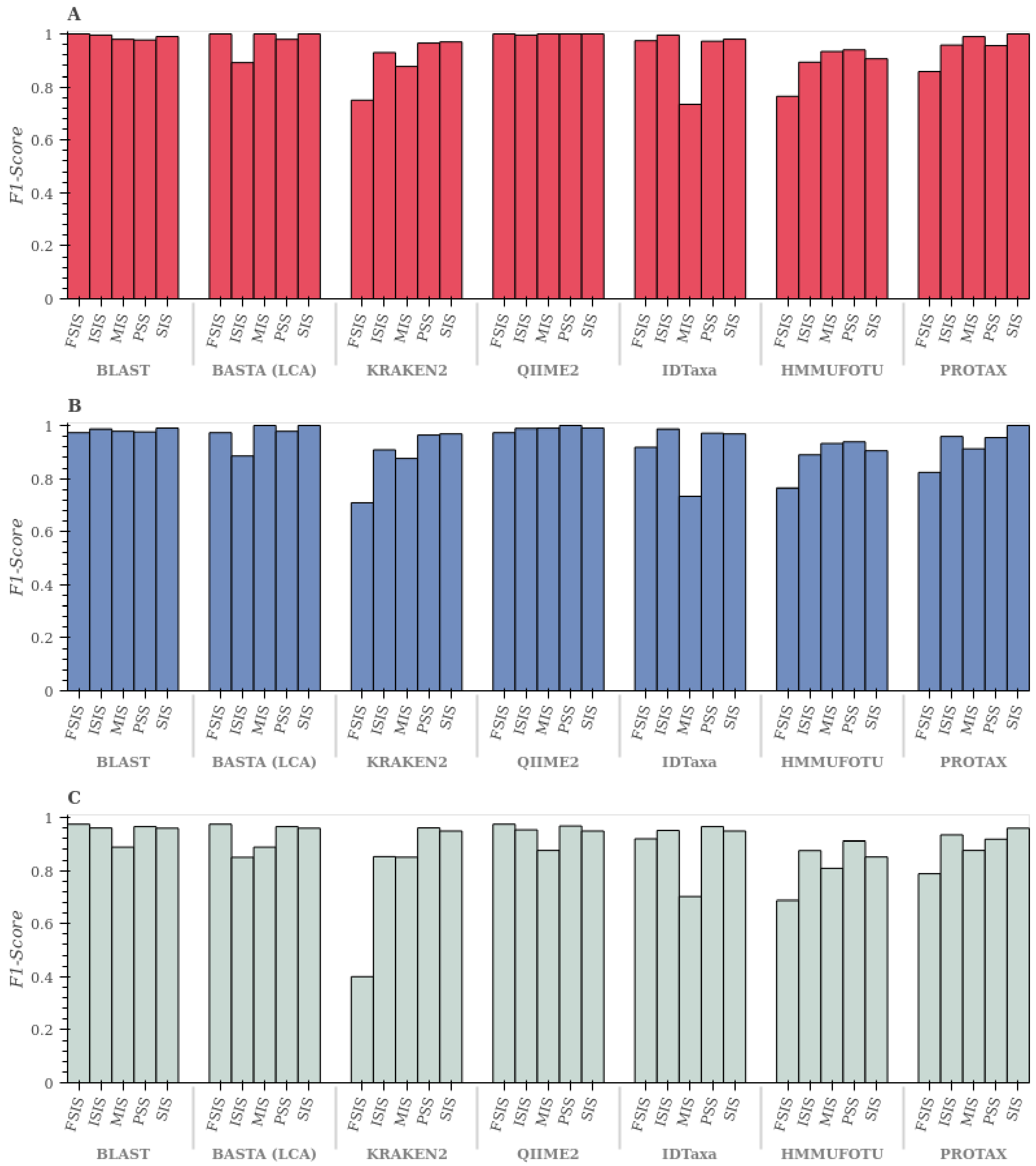

**Figure S2.** Box plot of the MCC for each of the mock communities.

FSIS: Fish single individual per species; ISIS: Insect single individual per species; MIS: Zooplankton multiple individuals per species; PSS: Zooplankton population of single species; SIS: Zooplankton single individual per species. MCC: Mathews correlation coefficient.

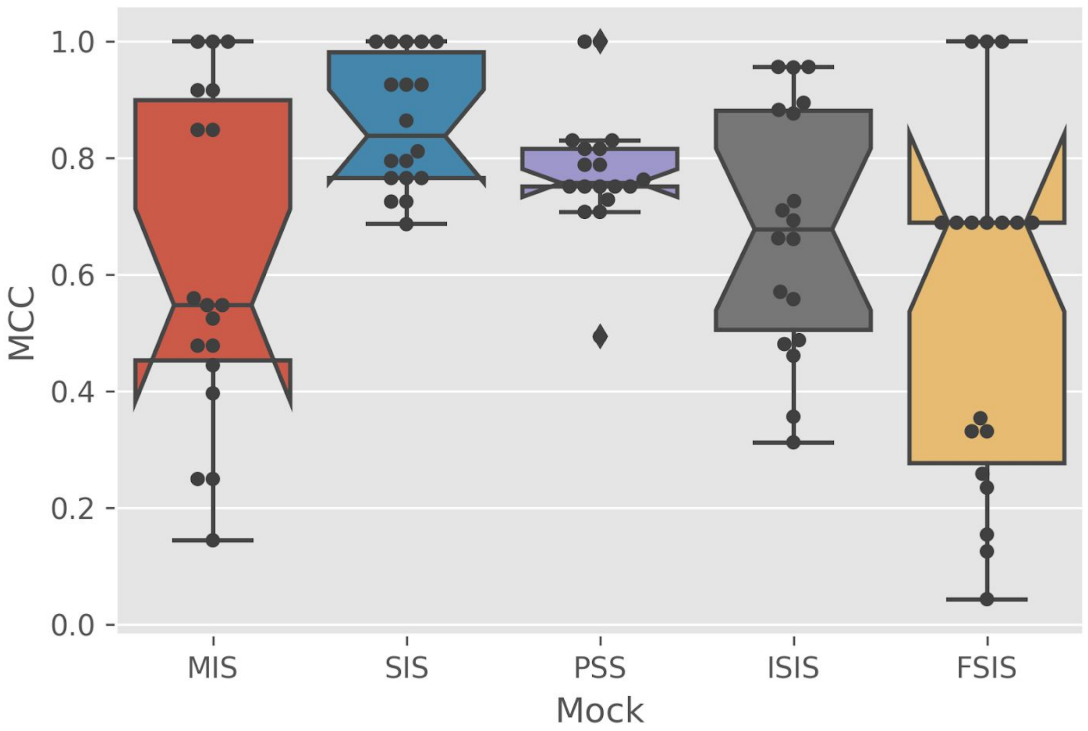

#### Supplementary Tables

**Table S1**. Fish Mock community composition.

| Order | Family | Genus | Species |
|---|---|---|---|
| Acipenseriformes | Acipenseridae | Acipenser | *Acipenser fulvescens* |
| Clupeiformes | Clupeidae | Alosa | *Alosa aestivalis* |
| Clupeiformes | Clupeidae | Alosa | *Alosa pseudoharengus* |
| Centrarchiformes | Centrarchidae | Ambloplites | *Ambloplites rupestris* |
| Siluriformes | Ictaluridae | Ameiurus | *Ameiurus nebulosus* |
| Anguilliformes | Anguillidae | Anguilla | *Anguilla rostrata* |
| Cypriniformes | Catostomidae | Catostomus | *Catostomus commersonii* |
| Salmoniformes | Salmonidae | Coregonus | *Coregonus artedi* |
| Perciformes | Gasterosteidae | Culaea | *Culaea inconstans* |
| Cypriniformes | Cyprinidae | Cyprinella | *Cyprinella spiloptera* |
| Cypriniformes | Cyprinidae | Cyprinus | *Cyprinus carpio* |
| Clupeiformes | Clupeidae | Dorosoma | *Dorosoma cepedianum* |
| Esociformes | Esocidae | Esox | *Esox lucius* |
| Esociformes | Esocidae | Esox | *Esox masquinongy* |
| Perciformes | Percidae | Etheostoma | *Etheostoma nigrum* |
| Perciformes | Percidae | Etheostoma | *Etheostoma flabellare* |
| Cyprinodontiformes | Fundulidae | Fundulus | *Fundulus diaphanus* |
| Hiodontiformes | Hiodontidae | Hiodon | *Hiodon tergisus* |
| Atheriniformes | Atherinopsidae | Labidesthes | *Labidesthes sicculus* |
| Gadiformes | Lotidae | Lota | *Lota lota* |
| Cypriniformes | Cyprinidae | Luxilus | *Luxilus cornutus* |
| Gadiformes | Gadidae | Microgadus | *Microgadus tomcod* |
| Centrarchiformes | Centrarchidae | Micropterus | *Micropterus dolomieu* |
| Perciformes | Moronidae | Morone | *Morone americana* |
| Perciformes | Moronidae | Morone | *Morone saxatilis* |
| Gobiiformes | Gobiidae | Neogobius | *Neogobius melanostomus* |
| Cypriniformes | Cyprinidae | Notemigonus | *Notemigonus crysoleucas* |
| Cypriniformes | Cyprinidae | Notropis | *Notropis atherinoides* |
| Cypriniformes | Cyprinidae | Notropis | *Notropis heterodon* |
| Cypriniformes | Cyprinidae | Notropis | *Notropis heterolepsis* |
| Cypriniformes | Cyprinidae | Notropis | *Notropis hudsonius* |
| Cypriniformes | Cyprinidae | Notropis | *Notropis rubellus* |
| Cypriniformes | Cyprinidae | Notropis | *Notropis volucellus* |
| Siluriformes | Ictaluridae | Noturus | *Noturus flavus* |

| | | | |
|---|---|---|---|
| Osmeriformes | Osmeridae | Osmerus | *Osmerus mordax* |
| Perciformes | Percidae | Perca | *Perca flavescens* |
| Perciformes | Percidae | Percina | *Percina caprodes* |
| Percopsiformes | Percopsidae | Percopsis | *Percopsis omiscomaycus* |
| Cypriniformes | Cyprinidae | Pimephales | *Pimephales promelas* |
| Salmoniformes | Salmonidae | Salmo | *Salmo trutta* |
| Perciformes | Percidae | Sander | *Sander vitreus* |

**Table S2.** Parameters used in cutadapt and DADA2 pipeline

| Program | Parameter | Value | Rationale |
|---|---|---|---|
| Cutadapt | -m | 170 | The minimum length parameter of the read before merging. Given that the Leray et al. (Leray *et al.* 2013) primers amplify a fragment of approximately 313 bp, each read must be at least 150 plus a 20bp overlap, hence 170 bp of minimum length filter |
| | -g/-G ; -a/-A | Primers | Removal of Forward (-g) and reverse (-G) primers. Given that amplification can occur on the reverse complement of the primers as well, we also filtered with the reverse complement (RC) of each primer with the options "-a" for the RC of the reverse primer and "-A" for the RC of the forward primer |
| | -u/-U | Estimated length | Trimming (-u) and reverse (-U) at the location previously recorded by the quality_plot function was passed to these options to trim the pair-end reads |
| | --match-read-wildcards | N/A | Because the Leray et al. (39) primers contain degenerate bases, this option is required for the proper adapter identification |
| | --trim-n | N/A | This option trimmed unknown/uncalled bases at the end of reads |
| | -n | 2 | To avoid chimeric primers within a read, we instructed cutadapt to search twice for the adapters |
| | --untrimmed-output / --untrimmed-paired-output | File names | Capture any untrimmed reads for a manual quality check |
| DADA2 | MaxEE | c(2, 2) | The maximum "expected errors" ("tolerance" during denoising) parameter for the reverse and forward reads |

| | | |
|---|---|---|
| truncQ | 2 | The truncation quality parameter |
| maxN | 0 | DADA2 pipeline does not accept unknown/uncalled bases in the read |
| pool | "pseudo" | Pool samples to denoise to use as priors in the pseudo-pooling approximation |
| minOverlap | 20 | Minimum number of base pairs to be overlapping for merging |
| matchIDs | TRUE | Match the sequence ID of forward and reverse for pairing |
| rm.phix | TRUE | Remove Illumina PhiX spikes if present |
| multithread | TRUE | Use multiple computational cpus where possible |

**Table S3.** Realized community in the mock after curation.

Shaded cells indicate the species originally intended. ASV size refers to the number of denoised sequences that correspond to a single amplicon sequence variant. ASV count refers to the number of ASVs detected. SIS: Single individual per species; MIS: Multiple individuals per species; PSS: Population of single species

TableS3

**Table S4**. Accuracy metrics used in benchmarking  directly or indirectly taxonomic assignment software.

NPV: Negative predictive value; MCC: Mathews correlation coefficient; AUC: Area under the curve.

| | Manuscript | | | | | | | | | | |
|---|---|---|---|---|---|---|---|---|---|---|---|
| **Metrics** | **Almeida et al. 2018** | **Austerlitz et al. 2009** | **Bazinet & Cummings 2012** | **Flynn et al. 2015** | **Gardner et al. 2019**** | **Hugerth & Anderson 2017** | **Lindgreen et al. 2017** | **McIntyre et al. 2017** | **Sczyrba et al. 2017** | **Siegwald et al. 2017** | **Total** |
| **Prevalence*** | 0 | 1 | 1 | 0 | 0 | 0 | 0 | 0 | 0 | 0 | 2 |
| **Recall** | 1 | 0 | 0 | 0 | 0 | 0 | 1 | 1 | 1 | 1 | 5 |
| **Precision** | 1 | 0 | 1 | 1 | 0 | 0 | 1 | 1 | 1 | 1 | 7 |
| **F1-Score** | 1 | 0 | 0 | 0 | 1 | 0 | 1 | 1 | 0 | 1 | 5 |
| **NPV** | 0 | 0 | 0 | 0 | 0 | 0 | 1 | 0 | 0 | 0 | 1 |
| **MCC** | 0 | 0 | 0 | 0 | 0 | 0 | 1 | 0 | 0 | 0 | 1 |
| **AUC** | 0 | 0 | 0 | 0 | 0 | 0 | 0 | 0 | 1 | 0 | 1 |
| **Base*** | 0 | 0 | 0 | 0 | 0 | 0 | 1 | 0 | 1 | 0 | 2 |
| **Chao** | 0 | 0 | 0 | 0 | 0 | 0 | 0 | 0 | 0 | 1 | 1 |
| **Bray** | 1 | 0 | 0 | 0 | 0 | 1 | 1 | 0 | 0 | 1 | 4 |
| **L1 norm** | 0 | 0 | 0 | 0 | 0 | 0 | 0 | 0 | 1 | 0 | 1 |
| **Unifrac** | 0 | 0 | 0 | 0 | 0 | 0 | 0 | 0 | 1 | 0 | 1 |
| **Fraction assigned reads** | 0 | 0 | 1 | 0 | 0 | 0 | 0 | 0 | 0 | 0 | 1 |

* It was called success rate by Austerlitz et al. 2009, but incorrectly called sensitivity and accuracy by Bazinet & Cummings 2012

** This is a meta-analysis that includes some other reported papers

*** Base metrics: This refers to the documentation of true and false positive and negative counts

**Table S5**. Description of programs selected for the taxonomic assignment benchmark

Table S5

**Table S6.** Performance and accuracy of the 6 programs evaluated.

TP: True positives; FP: False positives; FN: False negatives; FDR: False discovery rate; TPR: True positive rate; F1S: F1 Score; MCC: Mathews correlation coefficient; FSIS: Fish single individual per species; ISIS: Insect single individual per species; MIS: Zooplankton multiple individuals per species; PSS: Zooplankton population of single species; SIS: Zooplankton single individual per species.

| Mock | TP | FP | TN | FN | FDR | TPR | F1S | MCC | Shuff. | Total Calls | Time (s) | Memory (Mb) | CPU (%) | Taxonomic rank | Method |
|---|---|---|---|---|---|---|---|---|---|---|---|---|---|---|---|
| MIS | 48 | 2 | 6 | 0 | 0.04 | 1.00 | 0.98 | 0.85 | 6 | 56 | 0.75 | 423.88 | 17.71 | Family | |
| MIS | 48 | 2 | 6 | 0 | 0.04 | 1.00 | 0.98 | 0.85 | 6 | 56 | 0.75 | 423.88 | 17.71 | Genus | BLAST |
| MIS | 40 | 10 | 6 | 0 | 0.20 | 1.00 | 0.89 | 0.55 | 6 | 56 | 0.75 | 423.88 | 17.71 | Species | |
| MIS | 50 | 0 | 6 | 0 | 0.00 | 1.00 | 1.00 | 1.00 | 6 | 56 | 0.97 | 956.27 | 22.75 | Family | |
| MIS | 50 | 0 | 6 | 0 | 0.00 | 1.00 | 1.00 | 1.00 | 6 | 56 | 0.97 | 956.27 | 22.75 | Genus | LCA |
| MIS | 40 | 10 | 6 | 0 | 0.20 | 1.00 | 0.89 | 0.55 | 6 | 56 | 0.97 | 956.27 | 22.75 | Species | |
| MIS | 50 | 0 | 6 | 0 | 0.00 | 1.00 | 1.00 | 1.00 | 6 | 56 | 16.62 | 728.51 | 0.20 | Family | |
| MIS | 49 | 1 | 6 | 0 | 0.02 | 1.00 | 0.99 | 0.92 | 6 | 56 | 16.62 | 728.51 | 0.20 | Genus | QIIME |
| MIS | 39 | 11 | 6 | 0 | 0.22 | 1.00 | 0.88 | 0.52 | 6 | 56 | 16.62 | 728.51 | 0.20 | Species | |
| MIS | 39 | 2 | 6 | 9 | 0.05 | 0.81 | 0.88 | 0.44 | 6 | 56 | 0.46 | 728.30 | 0.39 | Family | |
| MIS | 39 | 1 | 6 | 10 | 0.03 | 0.80 | 0.88 | 0.48 | 6 | 56 | 0.46 | 728.30 | 0.39 | Genus | KRAKEN |
| MIS | 37 | 2 | 6 | 11 | 0.05 | 0.77 | 0.85 | 0.40 | 6 | 56 | 0.46 | 728.30 | 0.39 | Species | |
| MIS | 49 | 7 | 0 | 0 | 0.13 | 1.00 | 0.93 | -inf | 6 | 56 | 1.85 | 728.52 | 0.23 | Family | |
| MIS | 49 | 7 | 0 | 0 | 0.13 | 1.00 | 0.93 | -inf | 6 | 56 | 1.85 | 728.52 | 0.23 | Genus | HMMUFOTU |
| MIS | 38 | 18 | 0 | 0 | 0.32 | 1.00 | 0.81 | -inf | 6 | 56 | 1.85 | 728.52 | 0.23 | Species | |
| MIS | 29 | 2 | 6 | 19 | 0.06 | 0.60 | 0.73 | 0.25 | 6 | 56 | 8.09 | 728.51 | 0.20 | Family | |
| MIS | 29 | 2 | 6 | 19 | 0.06 | 0.60 | 0.73 | 0.25 | 6 | 56 | 8.09 | 728.51 | 0.20 | Genus | IDTAXA |
| MIS | 27 | 4 | 6 | 19 | 0.13 | 0.59 | 0.70 | 0.14 | 6 | 56 | 8.09 | 728.51 | 0.20 | Species | |
| MIS | 49 | 0 | 6 | 1 | 0.00 | 0.98 | 0.99 | 0.92 | 6 | 56 | 5.04 | 4217.98 | 57.70 | Family | |
| MIS | 42 | 7 | 6 | 1 | 0.14 | 0.98 | 0.91 | 0.56 | 6 | 56 | 5.04 | 4217.98 | 57.70 | Genus | PROTAX |
| MIS | 39 | 10 | 6 | 1 | 0.20 | 0.98 | 0.88 | 0.48 | 6 | 56 | 5.04 | 4217.98 | 57.70 | Species | |
| SIS | 50 | 1 | 7 | 0 | 0.02 | 1.00 | 0.99 | 0.93 | 7 | 58 | 1.03 | 4333.08 | 16.43 | Family | |
| SIS | 50 | 1 | 7 | 0 | 0.02 | 1.00 | 0.99 | 0.93 | 7 | 58 | 1.03 | 4333.08 | 16.43 | Genus | BLAST |
| SIS | 47 | 4 | 7 | 0 | 0.08 | 1.00 | 0.96 | 0.77 | 7 | 58 | 1.03 | 4333.08 | 16.43 | Species | |
| SIS | 51 | 0 | 7 | 0 | 0.00 | 1.00 | 1.00 | 1.00 | 7 | 58 | 1.19 | 1014.99 | 20.32 | Family | |
| SIS | 51 | 0 | 7 | 0 | 0.00 | 1.00 | 1.00 | 1.00 | 7 | 58 | 1.19 | 1014.99 | 20.32 | Genus | LCA |
| SIS | 47 | 4 | 7 | 0 | 0.08 | 1.00 | 0.96 | 0.77 | 7 | 58 | 1.19 | 1014.99 | 20.32 | Species | |
| SIS | 51 | 0 | 7 | 0 | 0.00 | 1.00 | 1.00 | 1.00 | 7 | 58 | 25.74 | 338.08 | 0.20 | Family | |
| SIS | 50 | 1 | 7 | 0 | 0.02 | 1.00 | 0.99 | 0.93 | 7 | 58 | 25.74 | 338.08 | 0.20 | Genus | QIIME |
| SIS | 46 | 5 | 7 | 0 | 0.10 | 1.00 | 0.95 | 0.73 | 7 | 58 | 25.74 | 338.08 | 0.20 | Species | |

| | | | | | | | | | | | | | | | |
|---|---|---|---|---|---|---|---|---|---|---|---|---|---|---|---|
| SIS | 48 | 1 | 7 | 2 | 0.02 | 0.96 | 0.97 | 0.80 | 7 | 58 | 0.46 | 818.87 | 10.73 | Family | |
| SIS | 48 | 1 | 7 | 2 | 0.02 | 0.96 | 0.97 | 0.80 | 7 | 58 | 0.46 | 818.87 | 10.73 | Genus | KRAKEN |
| SIS | 46 | 2 | 7 | 3 | 0.04 | 0.94 | 0.95 | 0.69 | 7 | 58 | 0.46 | 818.87 | 10.73 | Species | |
| SIS | 48 | 7 | 0 | 3 | 0.13 | 0.94 | 0.91 | -0.09 | 7 | 58 | 7.21 | 935.69 | 10.79 | Family | |
| SIS | 48 | 7 | 0 | 3 | 0.13 | 0.94 | 0.91 | -0.09 | 7 | 58 | 7.21 | 935.69 | 10.79 | Genus | HMMUFOTU |
| SIS | 43 | 12 | 0 | 3 | 0.22 | 0.93 | 0.85 | -0.12 | 7 | 58 | 7.21 | 935.69 | 10.79 | Species | |
| SIS | 49 | 2 | 7 | 0 | 0.04 | 1.00 | 0.98 | 0.86 | 7 | 58 | 12.96 | 686.05 | 4.39 | Family | |
| SIS | 48 | 3 | 7 | 0 | 0.06 | 1.00 | 0.97 | 0.81 | 7 | 58 | 12.96 | 686.05 | 4.39 | Genus | IDTAXA |
| SIS | 46 | 5 | 7 | 0 | 0.10 | 1.00 | 0.95 | 0.73 | 7 | 58 | 12.96 | 686.05 | 4.39 | Species | |
| SIS | 51 | 0 | 7 | 0 | 0.00 | 1.00 | 1.00 | 1.00 | 7 | 58 | 7.04 | 6715.03 | 60.55 | Family | |
| SIS | 51 | 0 | 7 | 0 | 0.00 | 1.00 | 1.00 | 1.00 | 7 | 58 | 7.04 | 6715.03 | 60.55 | Genus | PROTAX |
| SIS | 47 | 4 | 7 | 0 | 0.08 | 1.00 | 0.96 | 0.77 | 7 | 58 | 7.04 | 6715.03 | 60.55 | Species | |
| PSS | 213 | 10 | 23 | 0 | 0.04 | 1.00 | 0.98 | 0.82 | 23 | 246 | 4.27 | 7042.70 | 14.25 | Family | |
| PSS | 213 | 10 | 23 | 0 | 0.04 | 1.00 | 0.98 | 0.82 | 23 | 246 | 4.27 | 7042.70 | 14.25 | Genus | BLAST |
| PSS | 208 | 15 | 23 | 0 | 0.07 | 1.00 | 0.97 | 0.75 | 23 | 246 | 4.27 | 7042.70 | 14.25 | Species | |
| PSS | 214 | 9 | 23 | 0 | 0.04 | 1.00 | 0.98 | 0.83 | 23 | 246 | 4.48 | 1233.91 | 11.56 | Family | |
| PSS | 214 | 9 | 23 | 0 | 0.04 | 1.00 | 0.98 | 0.83 | 23 | 246 | 4.48 | 1233.91 | 11.56 | Genus | LCA |
| PSS | 208 | 15 | 23 | 0 | 0.07 | 1.00 | 0.97 | 0.75 | 23 | 246 | 4.48 | 1233.91 | 11.56 | Species | |
| PSS | 223 | 0 | 23 | 0 | 0.00 | 1.00 | 1.00 | 1.00 | 23 | 246 | 41.06 | 792.03 | 0.24 | Family | |
| PSS | 223 | 0 | 23 | 0 | 0.00 | 1.00 | 1.00 | 1.00 | 23 | 246 | 41.06 | 792.03 | 0.24 | Genus | QIIME |
| PSS | 209 | 14 | 23 | 0 | 0.06 | 1.00 | 0.97 | 0.76 | 23 | 246 | 41.06 | 792.03 | 0.24 | Species | |
| PSS | 208 | 0 | 23 | 15 | 0.00 | 0.93 | 0.97 | 0.75 | 23 | 246 | 0.54 | 790.04 | 0.88 | Family | |
| PSS | 208 | 0 | 23 | 15 | 0.00 | 0.93 | 0.97 | 0.75 | 23 | 246 | 0.54 | 790.04 | 0.88 | Genus | KRAKEN |
| PSS | 206 | 0 | 23 | 17 | 0.00 | 0.92 | 0.96 | 0.73 | 23 | 246 | 0.54 | 790.04 | 0.88 | Species | |
| PSS | 218 | 26 | 0 | 2 | 0.11 | 0.99 | 0.94 | -0.03 | 23 | 246 | 11.06 | 2002.83 | 40.64 | Family | |
| PSS | 218 | 26 | 0 | 2 | 0.11 | 0.99 | 0.94 | -0.03 | 23 | 246 | 11.06 | 2002.83 | 40.64 | Genus | HMMUFOTU |
| PSS | 206 | 38 | 0 | 2 | 0.16 | 0.99 | 0.91 | -0.04 | 23 | 246 | 11.06 | 2002.83 | 40.64 | Species | |
| PSS | 211 | 12 | 23 | 0 | 0.05 | 1.00 | 0.97 | 0.79 | 23 | 246 | 49.09 | 1169.00 | 4.01 | Family | |
| PSS | 211 | 12 | 23 | 0 | 0.05 | 1.00 | 0.97 | 0.79 | 23 | 246 | 49.09 | 1169.00 | 4.01 | Genus | IDTAXA |
| PSS | 208 | 15 | 23 | 0 | 0.07 | 1.00 | 0.97 | 0.75 | 23 | 246 | 49.09 | 1169.00 | 4.01 | Species | |
| PSS | 204 | 0 | 23 | 19 | 0.00 | 0.91 | 0.96 | 0.71 | 23 | 246 | 25.20 | 7122.03 | 69.26 | Family | |
| PSS | 204 | 0 | 23 | 19 | 0.00 | 0.91 | 0.96 | 0.71 | 23 | 246 | 25.20 | 7122.03 | 69.26 | Genus | PROTAX |
| PSS | 189 | 15 | 23 | 19 | 0.07 | 0.91 | 0.92 | 0.49 | 23 | 246 | 25.20 | 7122.03 | 69.26 | Species | |
| ISIS | 232 | 2 | 24 | 0 | 0.01 | 1.00 | 1.00 | 0.96 | 24 | 258 | 15.61 | 927.73 | 89.96 | Family | |
| ISIS | 228 | 6 | 24 | 0 | 0.03 | 1.00 | 0.99 | 0.88 | 24 | 258 | 15.61 | 927.73 | 89.96 | Genus | BLAST |
| ISIS | 216 | 18 | 24 | 0 | 0.08 | 1.00 | 0.96 | 0.73 | 24 | 258 | 15.61 | 927.73 | 89.96 | Species | |
| ISIS | 188 | 4 | 24 | 42 | 0.02 | 0.82 | 0.89 | 0.48 | 24 | 258 | 19.51 | 1626.13 | 5.85 | Family | |
| ISIS | 186 | 5 | 24 | 43 | 0.03 | 0.81 | 0.89 | 0.46 | 24 | 258 | 19.51 | 1626.13 | 5.85 | Genus | LCA |
| ISIS | 173 | 10 | 24 | 51 | 0.05 | 0.77 | 0.85 | 0.36 | 24 | 258 | 19.51 | 1626.13 | 5.85 | Species | |

| | | | | | | | | | | | | | | | |
|---|---|---|---|---|---|---|---|---|---|---|---|---|---|---|---|
| ISIS | 232 | 0 | 24 | 2 | 0.00 | 0.99 | 1.00 | 0.96 | 24 | 258 | 394.33 | 1420.47 | 0.24 | Family | |
| ISIS | 229 | 2 | 24 | 3 | 0.01 | 0.99 | 0.99 | 0.90 | 24 | 258 | 394.33 | 1420.47 | 0.24 | Genus | QIIME |
| ISIS | 213 | 16 | 24 | 5 | 0.07 | 0.98 | 0.95 | 0.66 | 24 | 258 | 394.33 | 1420.47 | 0.24 | Species | |
| ISIS | 203 | 5 | 24 | 26 | 0.02 | 0.89 | 0.93 | 0.57 | 24 | 258 | 0.76 | 1418.73 | 0.94 | Family | |
| ISIS | 195 | 9 | 24 | 30 | 0.04 | 0.87 | 0.91 | 0.49 | 24 | 258 | 0.76 | 1418.73 | 0.94 | Genus | KRAKEN |
| ISIS | 174 | 20 | 24 | 40 | 0.10 | 0.81 | 0.85 | 0.31 | 24 | 258 | 0.76 | 1418.73 | 0.94 | Species | |
| ISIS | 208 | 24 | 0 | 26 | 0.10 | 0.89 | 0.89 | -0.11 | 24 | 258 | 34.99 | 1419.10 | 0.23 | Family | |
| ISIS | 207 | 25 | 0 | 26 | 0.11 | 0.89 | 0.89 | -0.11 | 24 | 258 | 34.99 | 1419.10 | 0.23 | Genus | HMMUFOTU |
| ISIS | 201 | 31 | 0 | 26 | 0.13 | 0.89 | 0.88 | -0.12 | 24 | 258 | 34.99 | 1419.10 | 0.23 | Species | |
| ISIS | 232 | 1 | 24 | 1 | 0.00 | 1.00 | 1.00 | 0.96 | 24 | 258 | 569.28 | 1275.33 | 0.24 | Family | |
| ISIS | 228 | 4 | 24 | 2 | 0.02 | 0.99 | 0.99 | 0.88 | 24 | 258 | 569.28 | 1275.33 | 0.24 | Genus | IDTAXA |
| ISIS | 212 | 19 | 24 | 3 | 0.08 | 0.99 | 0.95 | 0.66 | 24 | 258 | 569.28 | 1275.33 | 0.24 | Species | |
| ISIS | 215 | 3 | 24 | 16 | 0.01 | 0.93 | 0.96 | 0.69 | 24 | 258 | 94.99 | 7557.84 | 22.24 | Family | |
| ISIS | 216 | 2 | 24 | 16 | 0.01 | 0.93 | 0.96 | 0.71 | 24 | 258 | 94.99 | 7557.84 | 22.24 | Genus | PROTAX |
| ISIS | 205 | 13 | 24 | 16 | 0.06 | 0.93 | 0.93 | 0.56 | 24 | 258 | 94.99 | 7557.84 | 22.24 | Species | |
| FSIS | 20 | 0 | 1 | 0 | 0.00 | 1.00 | 1.00 | 1.00 | 1 | 21 | 0.82 | 7541.88 | 1.70 | Family | |
| FSIS | 19 | 1 | 1 | 0 | 0.05 | 1.00 | 0.97 | 0.69 | 1 | 21 | 0.82 | 7541.88 | 1.70 | Genus | BLAST |
| FSIS | 19 | 1 | 1 | 0 | 0.05 | 1.00 | 0.97 | 0.69 | 1 | 21 | 0.82 | 7541.88 | 1.70 | Species | |
| FSIS | 20 | 0 | 1 | 0 | 0.00 | 1.00 | 1.00 | 1.00 | 1 | 21 | 1.02 | 1571.44 | 19.72 | Family | |
| FSIS | 19 | 1 | 1 | 0 | 0.05 | 1.00 | 0.97 | 0.69 | 1 | 21 | 1.02 | 1571.44 | 19.72 | Genus | LCA |
| FSIS | 19 | 1 | 1 | 0 | 0.05 | 1.00 | 0.97 | 0.69 | 1 | 21 | 1.02 | 1571.44 | 19.72 | Species | |
| FSIS | 20 | 0 | 1 | 0 | 0.00 | 1.00 | 1.00 | 1.00 | 1 | 21 | 20.35 | 1342.28 | 0.25 | Family | |
| FSIS | 19 | 1 | 1 | 0 | 0.05 | 1.00 | 0.97 | 0.69 | 1 | 21 | 20.35 | 1342.28 | 0.25 | Genus | QIIME |
| FSIS | 19 | 1 | 1 | 0 | 0.05 | 1.00 | 0.97 | 0.69 | 1 | 21 | 20.35 | 1342.28 | 0.25 | Species | |
| FSIS | 12 | 0 | 1 | 8 | 0.00 | 0.60 | 0.75 | 0.26 | 1 | 21 | 0.53 | 1341.75 | 0.62 | Family | |
| FSIS | 11 | 0 | 1 | 9 | 0.00 | 0.55 | 0.71 | 0.23 | 1 | 21 | 0.53 | 1341.75 | 0.62 | Genus | KRAKEN |
| FSIS | 5 | 0 | 1 | 15 | 0.00 | 0.25 | 0.40 | 0.13 | 1 | 21 | 0.53 | 1341.75 | 0.62 | Species | |
| FSIS | 13 | 1 | 0 | 7 | 0.07 | 0.65 | 0.76 | -0.16 | 1 | 21 | 6.40 | 1340.68 | 0.25 | Family | |
| FSIS | 13 | 1 | 0 | 7 | 0.07 | 0.65 | 0.76 | -0.16 | 1 | 21 | 6.40 | 1340.68 | 0.25 | Genus | HMMUFOTU |
| FSIS | 11 | 3 | 0 | 7 | 0.21 | 0.61 | 0.69 | -0.29 | 1 | 21 | 6.40 | 1340.68 | 0.25 | Species | |
| FSIS | 19 | 0 | 1 | 1 | 0.00 | 0.95 | 0.97 | 0.69 | 1 | 21 | 7.38 | 1340.72 | 0.25 | Family | |
| FSIS | 17 | 2 | 1 | 1 | 0.11 | 0.94 | 0.92 | 0.33 | 1 | 21 | 7.38 | 1340.72 | 0.25 | Genus | IDTAXA |
| FSIS | 17 | 2 | 1 | 1 | 0.11 | 0.94 | 0.92 | 0.33 | 1 | 21 | 7.38 | 1340.72 | 0.25 | Species | |
| FSIS | 15 | 0 | 1 | 5 | 0.00 | 0.75 | 0.86 | 0.35 | 1 | 21 | 2.63 | 7629.18 | 42.90 | Family | |
| FSIS | 14 | 1 | 1 | 5 | 0.07 | 0.74 | 0.82 | 0.15 | 1 | 21 | 2.63 | 7629.18 | 42.90 | Genus | PROTAX |
| FSIS | 13 | 2 | 1 | 5 | 0.13 | 0.72 | 0.79 | 0.04 | 1 | 21 | 2.63 | 7629.18 | 42.90 | Species | |

**Table S7**. Optimization values for the parameters for Blast.

TP: True positives; FP: False positives; FN: False negatives; FDR: False discovery rate; TPR: True positive rate; F1S: F1 Score; MCC: Mathews correlation coefficient; FSIS: Fish single individual per species; ISIS: Insect single individual per species; MIS: Zooplankton multiple individuals per species; PSS: Zooplankton population of single species; SIS: Zooplankton single individual per species.

Table S5

**Table S8a.** Optimization values for the parameters for LCA on the fish mock community (FSIS).

TP: True positives; FP: False positives; FN: False negatives; FDR: False discovery rate; TPR: True positive rate; F1S: F1 Score; MCC: Mathews correlation coefficient.

Table S6a

**Table S8b.** Optimization values for the parameters for LCA on the Insect mock community (ISIS).

TP: True positives; FP: False positives; FN: False negatives; FDR: False discovery rate; TPR: True positive rate; F1S: F1 Score; MCC: Mathews correlation coefficient.

Table S6b

**Table S8c.** Optimization values for the parameters for LCA on the zooplankton multiple individuals per species community (MIS).

TP: True positives; FP: False positives; FN: False negatives; FDR: False discovery rate; TPR: True positive rate; F1S: F1 Score; MCC: Mathews correlation coefficient.
Table S6c

**Table S8d**. Optimization values for the parameters for LCA on the zooplankton population single species (PSS) community.

TP: True positives; FP: False positives; FN: False negatives; FDR: False discovery rate; TPR: True positive rate; F1S: F1 Score; MCC: Mathews correlation coefficient.

Table S6d

**Table S8e.** Optimization values for the parameters for LCA on the zooplankton single individual per species mock community (SIS).

TP: True positives; FP: False positives; FN: False negatives; FDR: False discovery rate; TPR: True positive rate; F1S: F1 Score; MCC: Mathews correlation coefficient.

Table S6e

**Table S9**. Optimization values for the parameters for QIIME.

TP: True positives; FP: False positives; FN: False negatives; FDR: False discovery rate; TPR: True positive rate; F1S: F1 Score; MCC: Mathews correlation coefficient; FSIS: Fish single individual per species; ISIS: Insect single individual per species; MIS: Zooplankton multiple individuals per species; PSS: Zooplankton population of single species; SIS: Zooplankton single individual per species.

Table S9

**Table S10**. Optimization values for the parameters for Kraken.

TP: True positives; FP: False positives; FN: False negatives; FDR: False discovery rate; TPR: True positive rate; F1S: F1 Score; MCC: Mathews correlation coefficient; FSIS: Fish single individual per species; ISIS: Insect single individual per species; MIS: Zooplankton multiple individuals per species; PSS: Zooplankton population of single species; SIS: Zooplankton single individual per species.

Table S10

**Table S11**. Optimization values for the parameters for HMMUFOTU.

TP: True positives; FP: False positives; FN: False negatives; FDR: False discovery rate; TPR: True positive rate; F1S: F1 Score; MCC: Mathews correlation coefficient; FSIS: Fish single individual per species; ISIS: Insect single individual per species; MIS: Zooplankton multiple individuals per species; PSS: Zooplankton population of single species; SIS: Zooplankton single individual per species.

Table S11

**Table S12**. Optimization values for the parameters for IDtaxa.

TP: True positives; FP: False positives; FN: False negatives; FDR: False discovery rate; TPR: True positive rate; F1S: F1 Score; MCC: Mathews correlation coefficient; FSIS: Fish single individual per species; ISIS: Insect single individual per species; MIS: Zooplankton multiple individuals per species; PSS: Zooplankton population of single species; SIS: Zooplankton single individual per species.

Table S12

**Table S13**. Optimization values for the parameters for Protax.

TP: True positives; FP: False positives; FN: False negatives; FDR: False discovery rate; TPR: True positive rate; F1S: F1 Score; MCC: Mathews correlation coefficient; FSIS: Fish single individual per species; ISIS: Insect single individual per species; MIS: Zooplankton multiple individuals per species; PSS: Zooplankton population of single species; SIS: Zooplankton single individual per species.

Table S13

#### Supplementary methods

#### Methods S1

##### Summary of methods

Their source publications detail methodologies for DNA extraction and library preparation for the zooplankton (Zhang et al. 2018b), insect (Braukmann et al. 2019), and fish communities (next sections). Briefly, the zooplankton communities were assembled from DNA extracted from morphologically identified specimens using Qiagen DNeasy Blood & Tissue kits for single individuals (for the SIS mock community) and bulk per species (for the PSS and MIS mock communities) and amplified using the 18S V4 (Zhan et al. 2013), FC (Shokralla et al. 2015), Leray (Leray et al., 2013), and Folmer (Folmer et al. 1994) primers. Six SIS libraries, six MIS libraries, and 12 PPS libraries were indexed using Nextera® XT Index Kit (300bp) and sequence in a single 2x300bp Illumina MiSeq run (Zhang et al. 2018b). The insect mock community was collected in Malaise traps deployed near Cambridge, Ontario, Canada, and DNA was extracted from a single leg from each identified specimen following (Ivanova et al. 2006). C_LepFolF/C_LepFolR primer pair (Hernández-Triana et al. 2014) was used to amplify the COI fragment, and sequenced on an ABI 3730xl DNA Analyzer (Applied Biosystems, Foster City, California, USA) (Braukmann et al. 2019). Finally, the fish mock community was assembled from 41 North American fish species obtained from the Ministère des Forêts, de la Faune et des Parcs (Québec). Muscle or fin tissue was extracted using Qiagen Blood and Tissue kits and the COI fragment was amplified using the Leray primer pairs (Leray et al., 2013) and were sequenced using 2x300bp Illumina MiSeq with Nextera Libraries in a single run.

##### Sampling and identification of fish

Tissue extracts of 41 North American fish species were obtained from the Ministère des Forêts, de la Faune et des Parcs (Québec). The selected species were chosen to represent a diversity of families within the Actinopterygii. Tissues were stored in ethanol at -20C before extraction.

Muscle or fin tissue was extracted using Qiagen Blood and Tissue kits according to the manufacturer's instructions and equimolarised to 15ng/µl using the Quant-iT Picogreen dsDNA assay kit.

**Library preparation and next-generation sequencing (NGS)**

Equimolar amounts of DNA were combined to make two replicate mock communities. The mixture of mock community DNA was amplified in triplicate with the Leray primer with the following PCR chemistry: 7.875µl nuclease free water, 1.25µl 10X buffer (Genscript), 1mM $MgCl_2$ (ThermoFisher Scientific), 0.1mM GeneDirex dNTPs, 0.0125mg bovine serum albumen (ThermoFisher Scientific), 0.2mM each 10mM primer, 1.25U *taq* (GenScript) and 2µl DNA in a final volume of 12.5µl. The thermocycling regime followed that of the original paper. Amplicons were run on 1% agarose gel stained with SYBR™ Safe DNA Gel Stain (Thermo-Fisher Scientific) and visualized with UV light. Triplicate PCR amplicons for each sample were combined, cleaned with AMPure beads in a 0.75 ratio and indexed with the Nextera DNA indexing kit for 96 samples (Illumina). A second clean-up with AMPure beads was performed, and libraries were quantified and normalised to 5ng/ul. Sequencing was conducted using 2x300bp Illumina MiSeq at the McGill University and Génome Québec Innovation Centre, Montréal.

Leray, M., Yang, J.Y., Meyer, C.P. *et al.* A new versatile primer set targeting a short fragment of the mitochondrial COI region for metabarcoding metazoan diversity: application for characterizing coral reef fish gut contents. *Front Zool* **10,** 34 (2013). https://doi.org/10.1186/1742-9994-10-34

**Availability**

The sequence data will be available in Dryad upon manuscript acceptance

#### Supplementary notes

##### Supplementary Note S1

##### On the complexity and accuracy gain of the methods evaluated

It has been suggested that the increased complexity of methods has not substantially improved the accuracy of assignments (Bazinet & Cummings 2012; Garcia-Etxebarria *et al.* 2014), especially for higher taxonomic categories (e.g. genus and species). Disparities in the ways that taxonomic assignment software are compared, complicate the selection of the right tool (Escobar-Zepeda *et al.* 2018), especially for complex and diverse samples. Although benchmarking studies have been performed (Bazinet & Cummings 2012; Peabody *et al.* 2015; Lindgreen *et al.* 2016; Siegwald *et al.* 2017; McIntyre *et al.* 2017; Sczyrba *et al.* 2017; Almeida *et al.* 2018), there are significant disparities in their rankings of assignment software (Gardner *et al.* 2019) and they have not included the latest methods. Furthermore, potential bias arises when the performance of the method is evaluated by its developer (such as Peabody *et al.* 2015; McIntyre *et al.* 2017; Sczyrba *et al.* 2017; as exposed in Gardner *et al.* 2019). At a minimum, benchmarking studies should assess methods in an unbiased way, using real and *in silico* mock communities. Benchmarks which include more complex methods will be yet more informative to ecologists, but these have been infrequently addressed. Also, the compositional heterogeneity (i.e. variation in diversity and number of individuals per species) of real samples hasn't been taken into account, which can mislead the choice of the most appropriate methods. Finally, current benchmarking efforts tend to test the default parameters of the software, which might not be optimal for diverse communities. Most prior benchmarking studies have examined shotgun metagenomics, or have only been evaluated with mock communities that included fewer than ten species (Lu *et al.* 2017; Gardner *et al.* 2019) and therefore these results are unlikely to resemble ecological sampling.

**On parameter optimization**

Current benchmarking efforts tend to test only default parameters of a software which might not be optimal for diverse communities. Gardner et al. (2019) concluded that taxonomic assignments remain challenging, and stressed the importance of unbiased benchmarks (where the developers of the software are not involved in its own benchmarking; Boulesteix *et al.* 2013), especially in eDNA-based metabarcoding studies. Moreover, prior benchmarking studies (Bazinet & Cummings 2012; Peabody *et al.* 2015; Lindgreen *et al.* 2016; Siegwald *et al.* 2017; McIntyre *et al.* 2017; Sczyrba *et al.* 2017; Almeida *et al.* 2018; Gardner *et al.* 2019) did not include the probabilistic method, with its only implementation being PROTAX (Somervuo *et al.* 2016), or the recent implementations of evolutionary placement algorithms (Mirarab *et al.* 2012; Barbera *et al.* 2019).

**On the compositional heterogeneity**

Besides the need to evaluate newer software, there is a need for realistic testing, including variables important to ecologists. For example, Zhang et al.(2018) is the only study which tested the effect of varying numbers of individuals per species (hereafter referred to as compositional heterogeneity) on the success of the taxonomic assignment. However, their study focused on examining the impacts of multiple markers rather than benchmarking taxonomic assignment software. The effect of the composition of the sample in the taxonomic assignment is crucial to understand the weaknesses of taxonomic assignment software and important information for ecologists during software selection.

**On the Accuracy metrics**

To better understand the accuracy metrics used in this manuscript, consider the situation where 10 reads in a mock community with species A to I representing sequences 1 to 9 and the $10^{th}$ sequence have been reshuffled as explained above. A software was tested against this dataset and found that sequences 1 to 6 were correctly assigned to species A to F and therefore a TP equals to 6. Sequences 7 and 8 were wrongly assigned to A and B and therefore FP equals to 2. Sequence 9 wasn't assigned to any species so the FN is 1. Finally, the shuffled sequence got assigned the label F, making TN equal to 0 (TN is the number of shuffled sequences minus the ones predicted to be something). With these basic counts, the TPR would be 0.86, which means that 86% of the predictions were correct. The PPV will be 0.75, meaning that 75% of the label predictions were assigned to the right label. Overall, the F1-score of 0.8 indicates that the test is, on average, 80% accurate (here used semantically). The FDR was 0.25, and indicates that a false discovery was made 25% of the time. Finally, the MCC of -0.19 indicates that the method performed poorly with a strong tendency to wrongly predict labels.

##### On the practical context of the method's performance

In practical terms, during classification, all the programs evaluated here with the size of the mocks and databases used can be potentially run on a laptop, but the time frame could potentially be very long. IDtaxa, for example, could take up to an hour per classification with our datasets. However, if optimizing the parameters, multiple runs have to be made and therefore the scalability of the program becomes relevant. Programs like QIIME, BLAST, and KRAKEN2 are easily parallelizable to account for this, especially QIIME, but the memory requirements of programs like IDtaxa might be prohibitive in a commercial laptop and more suited for high performance computers (HPC or supercomputers), since the memory usage scales with the number of cpus used.

#### References


Almeida, A., Mitchell, A.L., Tarkowska, A. & Finn, R.D. (2018). Benchmarking taxonomic assignments based on 16S rRNA gene profiling of the microbiota from commonly sampled environments. *GigaScience*, **7**.

Barbera, P., Kozlov, A.M., Czech, L., Morel, B., Darriba, D., Flouri, T. & Stamatakis, A. (2019). EPA-ng: Massively Parallel Evolutionary Placement of Genetic Sequences. *Systematic Biology*, **68**, 365–369.

Bazinet, A.L. & Cummings, M.P. (2012). A comparative evaluation of sequence classification programs. *BMC Bioinformatics*, **13**, 92.

Boulesteix, A.-L., Lauer, S. & Eugster, M.J.A. (2013). A plea for neutral comparison studies in computational sciences. *Plos One*, **8**, e61562.

Escobar-Zepeda, A., Godoy-Lozano, E.E., Raggi, L., Segovia, L., Merino, E., Gutiérrez-Rios, R.M., Juarez, K., Licea-Navarro, A.F., Pardo-Lopez, L. & Sanchez-Flores, A. (2018). Analysis of sequencing strategies and tools for taxonomic annotation: Defining standards for progressive metagenomics. *Scientific Reports*, **8**, 12034.

Garcia-Etxebarria, K., Garcia-Garcerà, M. & Calafell, F. (2014). Consistency of metagenomic assignment programs in simulated and real data. *BMC Bioinformatics*, **15**, 90.

Gardner, P.P., Watson, R.J., Morgan, X.C., Draper, J.L., Finn, R.D., Morales, S.E. & Stott, M.B. (2019). Identifying accurate metagenome and amplicon software via a meta-analysis of sequence to taxonomy benchmarking studies. *PeerJ*, **7**, e6160.

Leray, M., Yang, J.Y., Meyer, C.P., Mills, S.C., Agudelo, N., Ranwez, V., Boehm, J.T. & Machida, R.J. (2013). A new versatile primer set targeting a short fragment of the mitochondrial COI region for metabarcoding metazoan diversity: application for characterizing coral reef fish gut contents. *Frontiers in zoology*, **10**, 34.

Lindgreen, S., Adair, K.L. & Gardner, P.P. (2016). An evaluation of the accuracy and speed of metagenome analysis tools. *Scientific Reports*, **6**, 19233.

Lu, J., Breitwieser, F.P., Thielen, P. & Salzberg, S.L. (2017). Bracken: estimating species abundance in metagenomics data. *PeerJ Computer Science*, **3**, e104.

McIntyre, A.B.R., Ounit, R., Afshinnekoo, E., Prill, R.J., Hénaff, E., Alexander, N., Minot, S.S., Danko, D., Foox, J., Ahsanuddin, S., Tighe, S., Hasan, N.A., Subramanian, P., Moffat, K., Levy, S., Lonardi, S., Greenfield, N., Colwell, R.R., Rosen, G.L. & Mason, C.E. (2017). Comprehensive benchmarking and ensemble approaches for metagenomic classifiers. *Genome Biology*, **18**, 182.

Mirarab, S., Nguyen, N. & Warnow, T. (2012). SEPP: SATé-enabled phylogenetic placement. *Pacific Symposium on Biocomputing*, 247–258.

Peabody, M.A., Van Rossum, T., Lo, R. & Brinkman, F.S.L. (2015). Evaluation of shotgun metagenomics sequence classification methods using in silico and in vitro simulated communities. *BMC Bioinformatics*, **16**, 363.

Sczyrba, A., Hofmann, P., Belmann, P., Koslicki, D., Janssen, S., Dröge, J., Gregor, I., Majda, S., Fiedler, J., Dahms, E., Bremges, A., Fritz, A., Garrido-Oter, R., Jørgensen, T.S., Shapiro, N., Blood, P.D., Gurevich, A., Bai, Y., Turaev, D., DeMaere, M.Z. & McHardy, A.C. (2017). Critical Assessment of Metagenome Interpretation-a benchmark of metagenomics software. *Nature Methods*, **14**, 1063–1071.

Siegwald, L., Touzet, H., Lemoine, Y., Hot, D., Audebert, C. & Caboche, S. (2017). Assessment of common and emerging bioinformatics pipelines for targeted metagenomics. *Plos One*, **12**, e0169563.

Somervuo, P., Koskela, S., Pennanen, J., Henrik Nilsson, R. & Ovaskainen, O. (2016). Unbiased probabilistic taxonomic classification for DNA barcoding. *Bioinformatics*, **32**, 2920–2927.

Zhang, G.K., Chain, F.J.J., Abbott, C.L. & Cristescu, M.E. (2018). Metabarcoding using multiplexed markers increases species detection in complex zooplankton communities. *Evolutionary applications*, **11**, 1901–1914.
